# Coordination of genome replication and anaphase entry by rDNA copy number in *S. cerevisiae*

**DOI:** 10.1101/2021.02.25.432950

**Authors:** Elizabeth X. Kwan, Gina M. Alvino, Kelsey L. Lynch, Paula F. Levan, Haley M. Amemiya, Xiaobin S. Wang, Sarah A. Johnson, Joseph C. Sanchez, Madison A. Miller, Mackenzie Croy, Seung-been Lee, Maria Naushab, Josh T. Cuperus, Bonita J. Brewer, Christine Queitsch, M. K. Raghuraman

**Author notes:** Further information and requests for resources and reagents should be directed to and will be fulfilled by M. K. Raghuraman.

## Abstract

Eukaryotes maintain hundreds of copies of ribosomal DNA (rDNA), many more than required for ribosome biogenesis, suggesting a yet undefined role for large rDNA arrays outside of ribosomal RNA synthesis. We demonstrate that reducing the *Saccharomyces cerevisiae* rDNA array to 35 copies, which is sufficient for ribosome function, shifts rDNA from being the latest replicating region in the genome to one of the earliest. This change in replication timing results in delayed genome-wide replication and classic replication defects. We present evidence that the requirement for rDNA to replicate late, which is conserved among eukaryotes, also coordinates the completion of genome replication with anaphase entry through the proper sequestration of the mitotic exit regulator Cdc14p in the rDNA-containing nucleolus. Our findings suggest that, instead of being a passive repetitive element, the large late-replicating rDNA array plays an active role in genome replication and cell cycle control.

## INTRODUCTION

Ribosomal DNA (rDNA) is uniquely situated at the intersection of ribosome biogenesis and genome replication, fundamental processes required for cell growth and proliferation. The rDNA sequence is comprised of four ribosomal RNA genes, which encode the core components of the ribosome; multiple rDNA copies are arranged in tandem to form large arrays. Many species have hundreds of rDNA copies per haploid genome: *e.g*., 90-300 in *Saccharomyces cerevisiae*, 70-400 in *Caenorhabditis elegans*, 80-600 in *Drosophila melanogaster*, 500-2500 in *Arabidopsis thaliana*, and 30-800 in humans (Mohan and Ritossa, 1970; Morton et al., 2020; Parks et al., 2018; Thompson et al., 2013). As the high rDNA copy number is often in substantial excess of what is required to meet ribosome demands, the majority of rDNA repeats are silenced (Conconi et al., 1992, 1989; Dammann et al., 1993; French et al., 2003; McStay and Grummt, 2008; Ye and Eickbush, 2006). Nevertheless, active maintenance of seemingly overabundant rDNA copies is conserved (Nelson et al., 2019), hinting that high rDNA copy number may have roles beyond ribosome production.

The phenotypic consequences of rDNA copy number variation span a broad range of cellular processes. While ribosome deficiencies result when rDNA copy number drops below a certain threshold (Delany et al., 1994; French et al., 2003; Ritossa and Atwood, 1966; Sanchez et al., 2017), other phenotypes result from rDNA copy number variants that are sufficient for ribosome biogenesis (Gibbons et al., 2014; Ide et al., 2010; Lu et al., 2018; Michel et al., 2005; Paredes et al., 2011; Picart-Picolo et al., 2020; Wang and Lemos, 2017; Xu et al., 2017).

Additionally, the plastic nature of the rDNA locus allows copy number fluctuations in the face of stresses such as nutrient availability (Aldrich and Maggert, 2015) or replication defects (Ide et al., 2007; Lynch et al., 2019; Salim et al., 2017; Sanchez et al., 2017). Because rDNA copy number changes often occur in response to various forms of replication stress, we sought to ask whether the reverse is also true: does rDNA copy number variation impact genome stability by influencing genome replication?

Given the vast length of the highly repetitive rDNA arrays, replication initiation must occur within the rDNA sequence, as documented in several species (Bénard et al., 1995; Brewer and Fangman, 1988; Coffman et al., 2006; Hyrien and Méchali, 1992; Little et al., 1993; Lópezestraño et al., 1998). Extensive characterization in the budding yeast *S. cerevisiae* has shown that each 9.1 kb repeat contains a potential origin of replication (rDNA Autonomously Replicating Sequence or rARS, (Brewer et al., 1992; Miller and Kowalski, 1993)). For a wild type yeast rDNA array of 150 copies, only 30-40 of the 150 replication origins are estimated to initiate replication (“fire”) within an S phase (Brewer and Fangman, 1988; Linskens and Huberman, 1988). Genome-wide origin firing is limited by the low abundance of initiation factors that promote the temporal staggering of origin activation (Collart et al., 2013; Lynch et al., 2019; Mantiero et al., 2011; Yoshida et al., 2013). Hyperactivation of rDNA replication diverts replication factors away from unique regions of the genome (Shyian et al., 2016; Yoshida et al., 2014) and leads to persistent underreplication of certain genomic regions (Foss et al., 2017). We reasoned that reduction of rDNA copy number from the wild type burden of 100-200 copies would alleviate the competition for limiting initiation factors at the other 300 replication origins across the genome (Nieduszynski et al., 2007).

Here we assess genome replication in isogenic yeast strains with rDNA arrays that are either wild type in size (100-180 copies) or minimal (35 copies). Contrary to our expectations, we discovered that the minimal rDNA array does not reduce competition with non-rDNA origins, but instead drastically advances rDNA replication time and increases the density of active rDNA origins. The large burst of early rDNA initiations from the minimal rDNA array causes a delay in replication in the rest of the genome. Furthermore, loss of the replication fork barrier gene *FOB1* sensitizes minimal rDNA strains to DNA damage and premature exit into anaphase, suggesting new roles for both Fob1p and rDNA in monitoring S phase progression. These findings show that the rDNA array is not merely a passive repetitive element during replication, but an active force that coordinates genome replication and cell cycle progression.

## RESULTS

### Strains with minimal rDNA arrays are not defective in ribosome production

We generated isogenic strains with a reduced rDNA array at the endogenous rDNA locus (Figure 1A) using the pRDN1-Hyg plasmid-based method (Chernoff et al., 1994; Kobayashi et al., 2001; Kwan et al., 2013). About 20-25% of the isolates with rDNA reductions showed increased ploidy (Figure S1A-B). Because maintenance of reduced rDNA requires the deletion of *FOB1* to prevent rDNA recombination and expansion, we included both *FOB1* and *fob1Δ* strains with wild type rDNA arrays as control strains. We also examined strains with weakened rDNA origins (rDNA^RM^) (Kwan et al., 2013), which we hypothesized would further reduce rDNA replication initiation events and favor initiation at origins outside the rDNA locus. We generated strains with 35, 45, and 55 copies of rDNA and decided to focus on strains with the smallest rDNA array (minimal rDNA strains, 35 rDNA *fob1Δ* and 35 rDNA^RM^ *fob1Δ*) in comparison and compared these strains to those with wild type copy number (170 rDNA, 180 rDNA *fob1Δ*, and 100 rDNA^RM^).

**Figure 1.**
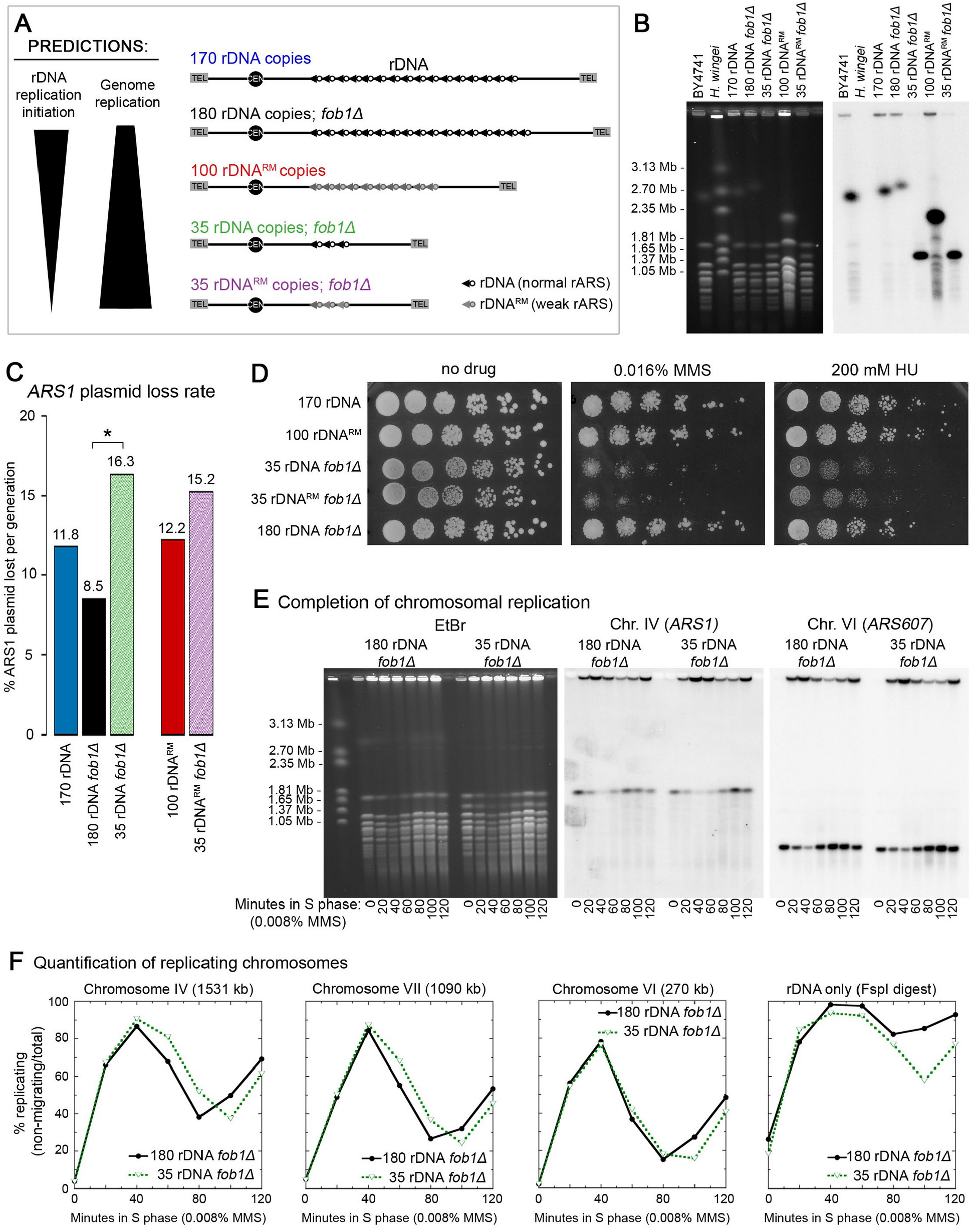
Strains with minimal rDNA copy number exhibit non-rDNA replication defects. (A) Depiction of rDNA arrays examined in this study and prediction of genome replication effects resulting from competition for replication factors. Strains differed at the rDNA locus in copy number and/or presence of a weak rDNA origin (rDNA^RM^). (B) CHEF gel confirmation of rDNA copy number. Ethidium bromide stained gel (left) and the resulting Southern blot (right) hybridized with a single copy Chr. XII probe (*CDC45*). (C) Loss of maintenance of an *ARS1*-containing plasmid was assessed in exponentially grown cultures. (*) indicates significant difference in plasmid loss rate calculated from slope variance (p = 0.03). (D) Spot assays for sensitivity to 200 mM HU and 0.016% MMS. (E) Migration assay of chromosomes from cells released into S phase in the presence of 0.08% MMS using CHEF gel electrophoresis and Southern blotting. Chromosomes are unable to migrate out of the plug/well while replicating (Hennessy & Botstein 1991), allowing for quantification of replicating vs. non-replicating chromosomes (fully replicated and unreplicated)by comparing the signal in the gel body vs. plug/well. Hybridization to specific sequences allows examination of individual chromosomes, IV and VI shown. (F) Quantification of the percentage of chromosomes undergoing replication in 0.008% MMS: chromosomes VI, IV, and VII as well as the isolated rDNA array. Completion of chromosome replication is reflected in the decrease of signal in the CHEF gel well.

Although previous work demonstrated that 35 rDNA copies are sufficient for wild type levels of rRNA synthesis and ribosome biogenesis (Dauban et al., 2019; French et al., 2003; Ide et al., 2010; Kim et al., 2006), we wanted to confirm the absence of ribosome biogenesis defects in the minimal rDNA strains. We assessed phenotypes due to ribosome insufficiency: slower growth rate, increased sensitivity to cycloheximide over a range of concentrations, and decreased relative 25S rRNA abundance (Abovich et al., 1985; Rosado et al., 2007; Sanchez et al., 2017). The minimal rDNA strains behaved similarly to the wild type rDNA controls in these assays (Figure S1C-E), confirming that rDNA reduction to 35 rDNA copies does not generate significant ribosome biogenesis defects.

### Minimal rDNA strains show reduced plasmid maintenance and increased sensitivity to MMS and hydroxyurea

We examined the effects of rDNA copy number reduction on plasmid maintenance, a basic assay that identifies mutants with DNA replication defects on the basis of their poor ability to replicate plasmids (Maine et al., 1984; Shima et al., 2007; Tye, 1999). Because replication initiation at a plasmid origin is subject to the same competition for replication factors as genomic origins, we expected that strains with minimal rDNA arrays would show improved plasmid maintenance compared to the wild type rDNA controls. However, both minimal rDNA strains displayed high loss rates of the *ARS1* (Autonomously Replicating Sequence 1, (Stinchcomb et al., 1979)) test plasmid: 16.3%/generation for the minimal rDNA strain and 15.2%/generation for the minimal rDNA^RM^ strain, almost double that of the control *fob1Δ* strain with wild type rDNA (8.5%/generation, *p* = 0.03) and higher than either of the wild type rDNA *FOB1* strains (11.8%/generation and 12.2%/generation, Figure 1C). Since the Southern blot method reflects the amount of plasmid relative to genomic DNA sequences in the culture (Brewer and Fangman, 1994), the observed differences can be ascribed to differences in plasmid replication rather than segregation. Counter to our prediction that rDNA reduction would improve DNA replication, the strains with minimal rDNA arrays instead showed defects in plasmid replication.

If minimal rDNA arrays are associated with defects in DNA replication, the strains carrying these arrays should be hypersensitive to conditions that induce replication stress and DNA damage (Shimada et al., 2002; Trabold et al., 2005). Hydroxyurea (HU) induces replication stress by inhibiting the production of dNTPs, which does not directly damage DNA (Alvino et al., 2007) but slows cell cycle progression through activation of the replication checkpoint (Santocanale and Diffley, 1998). In contrast, MMS is an alkylating agent that induces DNA damage (Paulovich and Hartwell, 1995). Consistent with previous work (Ide et al., 2010), strains with minimal rDNA showed greater sensitivity to DNA damage by MMS (Figure 1D). We found that rDNA reduction also conferred greater sensitivity to HU, suggesting that reducing rDNA copy number induces problems with both DNA replication and repair.

### A minimal rDNA strain shows delayed replication completion across multiple chromosomes

To determine if rDNA reduction leads to chromosomal replication defects, we performed an S phase specific CHEF gel electrophoresis assay on strains with wild type or minimal rDNA copy number. In CHEF gel electrophoresis, chromosomes that are undergoing replication will not migrate from the well (Hennessy et al., 1991; Lynch et al., 2019), while non-replicating chromosomes, either those that are pre-replication or have completed replication, migrate into the gel at their expected sizes. To measure replication completion for individual chromosomes, we quantified the relative amount of Southern blot hybridization signal remaining in the well at 20-minute intervals across a synchronous S phase. For this assay, G1 synchronized cells were released into S phase (Ide et al., 2010) and cell cycle progression was confirmed by flow cytometry (Figure S2A-B).

We examined replication completion for several chromosomes of different lengths as well as the rDNA array itself in *fob1Δ* strains with wild type or minimal rDNA. As expected, for samples collected in G1 prior to the onset of S phase, chromosomes migrated at their typical positions in the CHEF gel (0 minute samples, Figure 1E). After release into S phase, both strains began showing well-retention of chromosomes at the same time, indicating that the minimal rDNA strain does not have a delayed start of chromosomal replication. However, the two strains differed in their time of replication completion: Chromosomes IV (1531 kb) and VII (1090 kb) were delayed by 10-15 minutes in the 35 rDNA *fob1Δ* strain (Figure 1F, S1F). The extent of delay in replication completion appears to be proportional to chromosome length. We observed a smaller delay in replication completion for Chromosome II (813 kb) and barely any delay for chromosome VI (270 kb), the second smallest *S. cerevisiae* chromosome (Figure 1F, S1F-G).

We had difficulty quantifying Chromosome XII (containing the rDNA locus) for the wild type rDNA strain: 56% of chromosome XII did not migrate into the gel, likely due to its large size (Figure S1F, H). We instead quantified replication completion of intact rDNA arrays, using a digest to separate the rDNA locus from its chromosomal context (Figure S1F). We found that a higher fraction of the minimal rDNA array completed replication than the wild type rDNA array (Figure 1F). We conclude that the minimal rDNA strain shows significant genome-wide delays in completing chromosomal replication but fewer problems with completion of rDNA replication.

### Minimal rDNA strains advance rDNA replication and delay genome-wide replication

To investigate the kinetics of genome-wide replication in strains with reduced rDNA copy number, we performed density transfer experiments. Density transfer experiments exploit the semi-conservative nature of DNA replication to distinguish newly replicated DNA from unreplicated DNA at defined loci in the genome. Cells are grown in isotopically dense (“heavy”) medium, synchronized, and released into isotopically normal (“light”) medium such that the newly replicated DNA forms a “heavy/light” hybrid molecule. Hybrid replicated DNA can be resolved from unreplicated dense DNA by cesium chloride gradient fractionation and analyzed using quantitative Southern blotting and microarrays (Alvino et al., 2007; Raghuraman et al., 2001). Although this method cannot distinguish active origin firing from passive DNA replication, it allows us to identify replication differences across the genome. We collected samples across an S phase and assessed replication kinetics both on an individual locus level and on a genome-wide scale, confirming with flow cytometry that the minimal and wild type rDNA strains showed similar G1 arrest and progression through S phase (Figure S2C).

We initially focused on the rDNA locus, which is late-replicating in *S. cerevisiae* and several metazoan eukaryotic species (Coffman et al., 2005; Concia et al., 2018; Foss et al., 2017; Labit et al., 2008; Schübeler et al., 2002). We calculated rDNA T_rep_, the time at which half-maximal replication was achieved, for each strain. While the rDNA locus was indeed late-replicating in the wild type strain (T_rep_ =40.0’), the minimal rDNA array replicated earlier with a T_rep_ of 28.9’ (Figure 2A-B, S3A). In fact, the minimal rDNA locus was one of the earliest loci to replicate rather than one of the latest (Figure 2B). The minimal rDNA^RM^ array showed a similar advancement of replication time (Figure S3B), indicating that minimal rDNA arrays replicate early regardless of the rARS allele.

**Figure 2:**
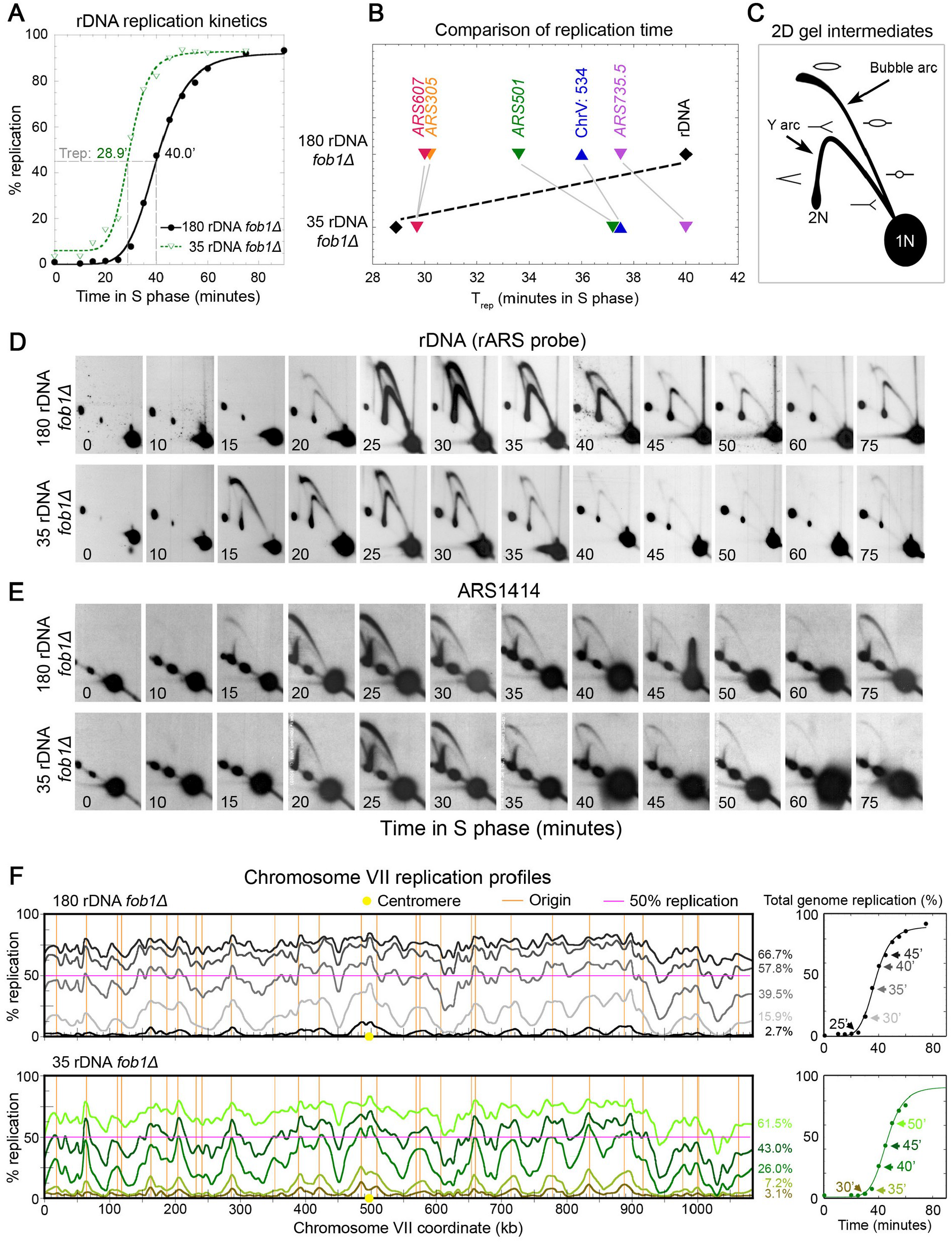
A minimal rDNA array replicates early and delays genome replication. We calculated rDNA T_rep_, the time at which half-maximal replication was achieved, for several loci. (A) The minimal rDNA array replicates 11 minutes earlier than the 180-copy rDNA array. Replication kinetic curves were generated from density transfer slot blot analysis. (B) Index of T_rep_ for different replication origins and genomic region Chr. V: 534. (C) Diagram of a 2D gel showing locations of actively replicating bubble or passively replicating Y intermediates. (D,E) 2D gel analysis of the rDNA origin of replication (rARS) and *ARS1414* throughout S phase. (F) Replicating DNA from multiple density transfer samples representing different time points were hybridized to microarrays and used to generate genome-wide replication profiles. Chromosome VII replication profiles across S phase for the 180 rDNA *fob1Δ* strain (top) and the 35 rDNA *fob1Δ* strain (bottom). Total genomic replication levels and time interval are indicated on the right. Locations of centromere and origins of replication are noted by yellow circles or orange lines. 50% replication threshold is denoted by pink horizontal line.

Because this drastic shift of rDNA replication timing from late to early might affect the order in which replication factors are recruited to other genomic origins, we compared the replication kinetics for several other genomic loci (Figure 2B, S3C-D, Table S2). *ARS305* and *ARS607*, two of the earliest and most efficient replication origins in *S. cerevisiae* (Friedman et al., 1997; Hoang et al., 2007), did not differ greatly in timing between the minimal and wild type rDNA strains. However, the late-replicating loci *ARS501* (also called *ARS522*), *ARS735.5*, and ChrV:534000 without an origin (Ferguson and Fangman, 1992) were delayed in the minimal rDNA strain, while their relative replication order was maintained.

### Altered replication timing is due to altered time of origin initiation

The observed shifts of replication timing could be the result of changes to the time of origin activation and/or efficiency. Although the rDNA kinetic curves (Figure 2A) suggested that the minimal rDNA array began replicating earlier than the wild type array, we wanted to directly examine replication initiation using 2D gel electrophoresis of synchronized cells sampled across S phase. If the time of replication initiation was altered, we should observe differences in when the replication “bubble” arc becomes visible among our strains (Figure 2C). Strong rDNA origin (rARS) initiation in the minimal rDNA locus began robustly at 15 minutes into S phase, whereas the wild type rDNA locus initiated replication later (at 20 minutes) (Figure 2D). These results confirm that early rDNA origin initiation is responsible for the earlier replication of the minimal rDNA locus seen by density transfer.

The altered replication timing of other genomic loci was also confirmed by 2D gels. In the minimal rDNA strain, we observed ∼5-minute delays in replication initiation at late origins such as *ARS1414* and *ARS735.5* (Figure 2E, S3E), with no apparent change in cumulative origin use. We found no difference in initiation timing at the early, efficient *ARS305* (Figure S3F), which reflects the density transfer data and is consistent with both the minimal rDNA and wild type strains entering S phase at the same time. We conclude that the changes to replication timing in the minimal rDNA strain are due to altered time of replication initiation at the rDNA and at late non-rDNA origins of replication.

### Delayed replication progression is seen genome-wide in the minimal rDNA strains

Since individual loci exhibited altered replication timing, we wanted to examine the effects genome-wide in strains with early-replicating minimal rDNA arrays. We hybridized the fractionated density transfer samples to microarrays and plotted percent replication against chromosomal coordinates; these plots create chromosomal profiles that describe the landscape of replication over time (Alvino et al., 2007; Raghuraman et al., 2001). We picked the earliest S phase sample in which replication was detected and the three subsequent 5-minute intervals. In the earliest sample with detectable hybrid density DNA at replication origins, we found only subtle differences between the replication profiles of the minimal rDNA strain and that of the wild type strain. (3.1% vs. 2.7% genome replicated, Figure 2F, S4, S5A-B). However, at the next 5 minute interval, the minimal rDNA strain showed lower levels of replication completion (7.2% genome replicated) compared to the wild type strain (15.9%) (Figure S5A). This trend continued for the subsequent intervals, consistent with slower advancement of replication across the genome for the minimal rDNA strain. We did not identify any specific genomic regions that were preferentially altered in their replication profile; chromosome-wide replication profiles from the minimal rDNA strain and wild type rDNA strain are superimposable when they achieve similar levels of genome replication (Figure S5B). These data, together with the single locus data, demonstrate that the minimal rDNA strain exhibits genome-wide replication delay without region-specific effects.

### Cyclin regulation reflects DNA replication delays in cells with minimal rDNA

Delayed completion of genome replication should manifest as delayed appearance of cell cycle landmarks. Such landmarks include the degradation of the CDK activator Clb5p (Schwob and Nasmyth, 1993), the phosphorylation of limiting initiation factor Sld2p (Bloom and Cross, 2007; Lynch et al., 2019; Masumoto et al., 2002), and the reinstatement of the CDK inhibitor Sic1p (Barberis et al., 2012; Khmelinskii et al., 2007; Schwob et al., 1994; Verma et al., 1997). We separately tagged Clb5p, Sld2p, and Sic1p at their endogenous loci using a 3HA tag (Longtine et al., 1998) in both the minimal rDNA and wild type rDNA strains. We then collected synchronized samples for each strain throughout S phase for analysis via flow cytometry and western blotting. The tagged minimal rDNA and wild type rDNA strains showed similar S phase progression (Figure S2E). However, for cyclin signaling, the minimal rDNA strain showed ∼10-minute delays in the degradation of Clb5p-HA, the phosphorylation of Sld2p-HA, and the degradation and reappearance of Sic1p-HA (Figure S3A-C). These delays in cyclin signaling in the minimal rDNA strain match the strain’s delayed genome-wide replication.

### How many rDNA origins in the minimal rDNA array initiate replication during early S phase?

Approximately one in five of the rDNA origins is thought to serve as a replication initiation site in cells with wild type rDNA (Brewer and Fangman, 1988; Linskens and Huberman, 1988). By this estimate, a strain with 180 rDNA copies would have ∼36 active origins in the rDNA locus—close to the maximum number of possible rDNA origin initiations in the minimal rDNA strains. We wondered how many rDNA origins fire in early S phase in the minimal rDNA strain and if increased early rDNA firing would suffice to alter replication timing genome-wide.

Digestion of replicating rDNA by the restriction enzyme NheI generates a variety of distinct molecular intermediates/structures such as “bubbles” from active initiation, passively replicated “Y” fragments, and “X” fragments produced by converging replication forks (Figure 4A), all of which can be resolved by 2D gel electrophoresis and Southern blotting (Figure 4B) (Brewer and Fangman, 1988, 1987). However, accurate quantification of early rDNA initiation requires the presence of the ribonucleotide reductase inhibitor hydroxyurea (HU) to slow down fast-moving replication forks; these forks in *S. cerevisiae* travel at a rate of 1.5 kb per minute at 30°C (Bell and Labib, 2016). Replication forks initiated at an rDNA origin will travel off the 4.7 kb origin-containing fragment (Figure 4A) in under 2 minutes and may fuse with oncoming rDNA forks in ∼3 minutes. We therefore released G1-synchronized cultures into S phase in the presence of HU, which allowed us to capture more replication intermediates and limit initiation events to early S phase (Alvino et al., 2007; Feng et al., 2006). In addition to the 4.7 kb rARS fragment from every rDNA repeat, the NheI digest also generates a 24.4 kb single-copy rARS fragment at the telomere proximal end of the rDNA array (Figure 4A-B). We used this single-copy rDNA fragment for normalization, allowing us to generate a “per cell” estimate of active rARS initiation in early S phase.

**Figure 3:**
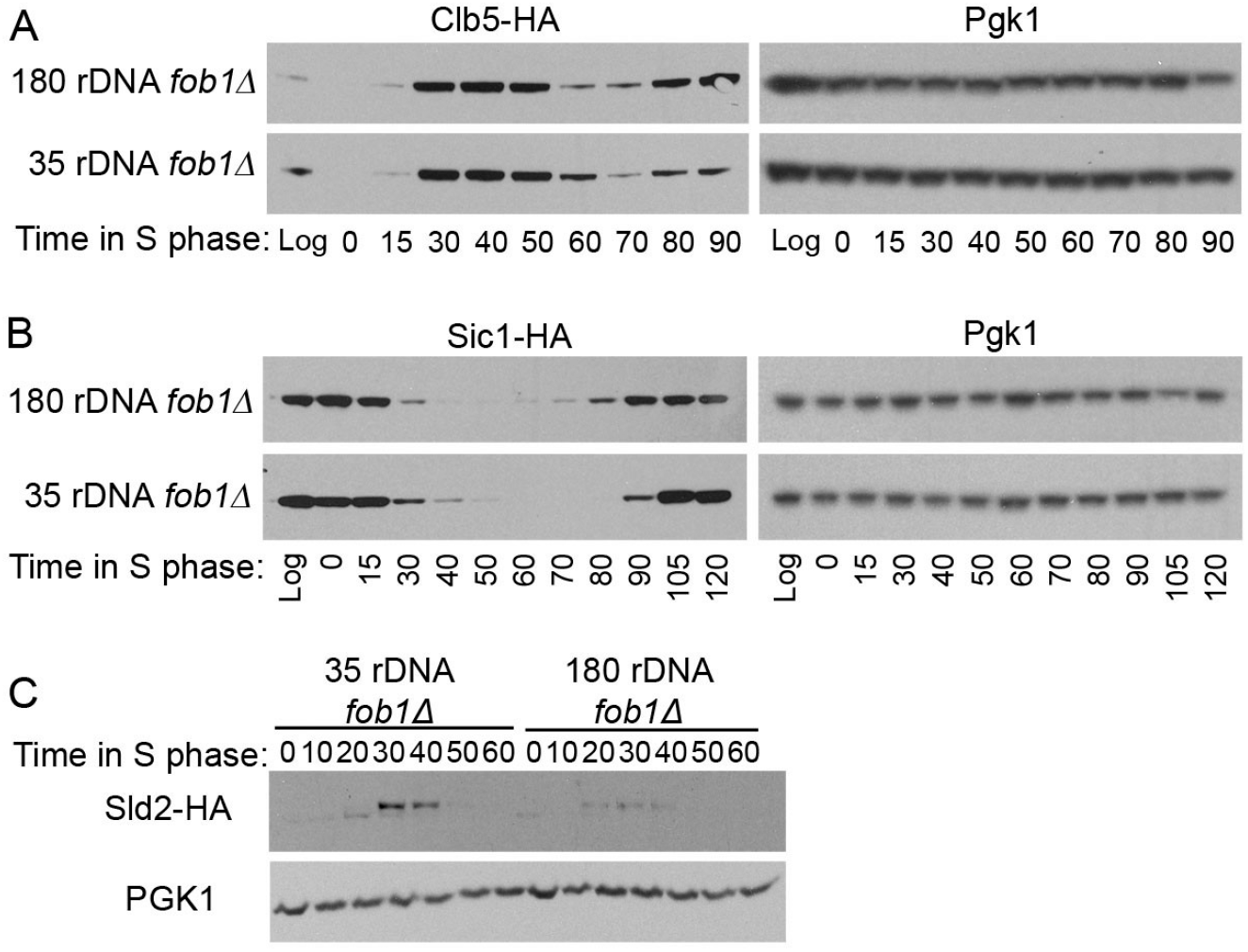
Cyclin regulation is delayed in minimal rDNA strains. Western blots of (A) HA-tagged Clb5, (B) HA-tagged Sic1, and (C) phosphorylation of HA-tagged Sld3 in synchronized cells progressing through S phase. Antibodies against Pgk1 were used to confirm similarly loaded protein concentration.

**Figure 4:**
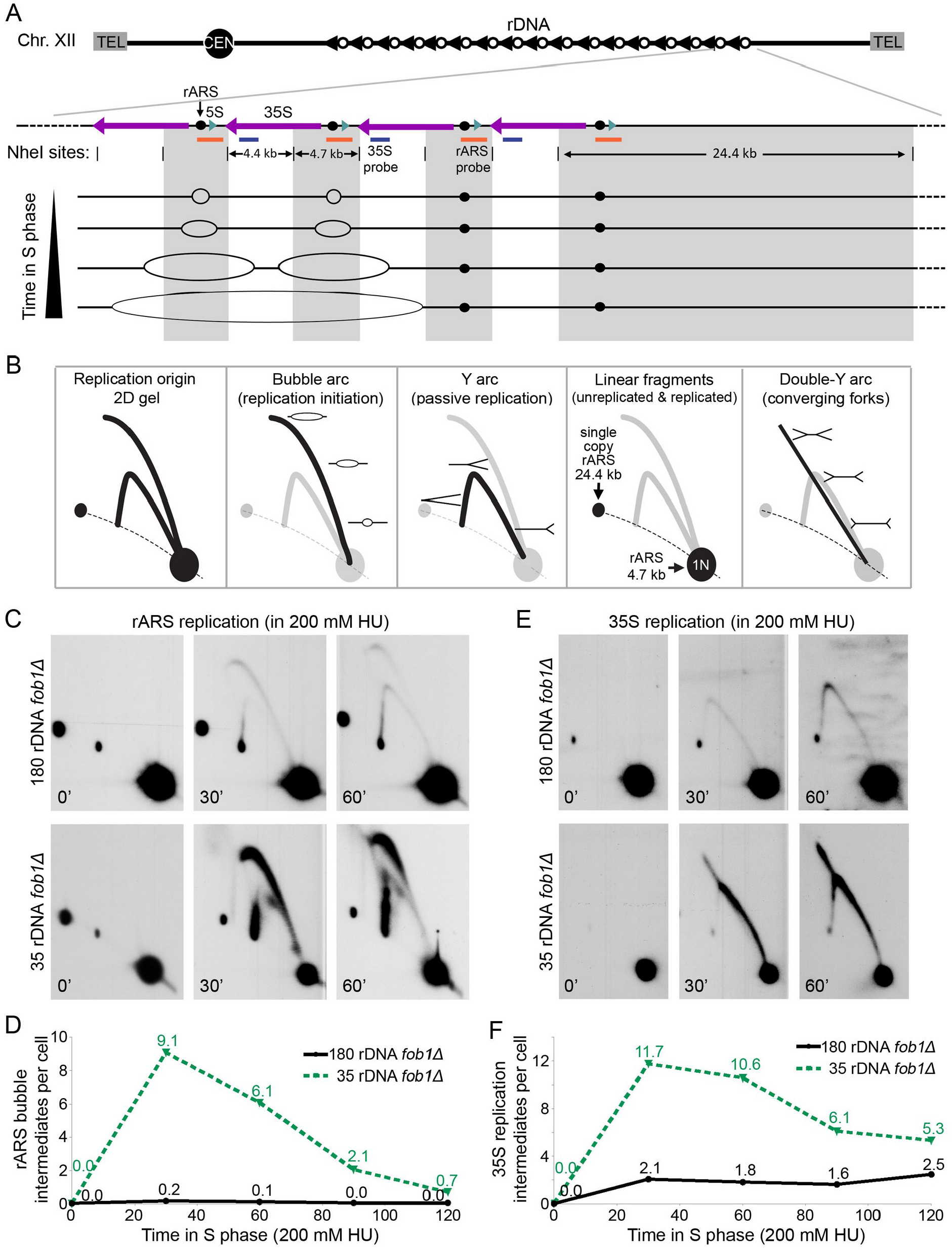
rDNA copy number reduction drives a 10-fold increase in rDNA replication initiation during early S phase. (A) Schematic of rDNA locus organization, relevant NheI-digest fragments, Southern blot probe locations, and replication intermediates predicted throughout S phase. The 24.4 kb fragment on the telomere proximal edge of the rDNA array serves as an internal reference for quantification as it is present in a single copy per cell and hybridizes to the rARS probe. (B) Representation of different replication intermediates on a 2D gel. (C,E) Timed 2D gels of cells released into S phase in the presence of 200 mM HU. 2D gels were probed with both (C) the rARS probe fragment and then (E) the 35S probe. Converging replication fork intermediates indicate initiation from adjacent rDNA repeats and are seen in the strain with 35 rDNA copies but not the strain with 180 rDNA copies. (D) Replication bubbles created by rARS initiations were quantified and normalized to the signal in the single copy 24.4 kb linear spot. (F) Estimation of replication fork intermediates present in the 4.4kb NheI fragment without the rARS. Total replication fork signal was normalized to the 4.4 kb 1N spot and adjusted for rDNA copy number.

Compared to the wild type rDNA strain, the minimal rDNA strain showed a far stronger rDNA bubble arc signal, which represents active replication initiations during S phase in HU (Figure 4C, S5C). At the 30 minute time point, we estimate that the minimal rDNA strain had 9.1 rDNA initiations per cell whereas the wild type rDNA strain had less than one (Figure 4D). This quantification is likely an underestimate due to the movement of replication forks off the rARS-containing fragment (Figure 4A). Passively replicating Y-arc and convergent double-Y intermediates (Figure 4B) seen in the adjacent non-origin 35S fragment (Figure 4E, S5D) represent replication forks that have traveled from a neighboring rDNA origin. Quantification of these Y and double-Y fragments relative to the linear 1N spot (Figure 4B) indicate that the wild type rDNA array had an additional 2 initiation events per cell while the minimal rDNA array had an additional 11 (Figure 4F). Combining active and passive replication values, we estimate that the wild type strain contains 2 active rDNA origins in early S phase while the minimal rDNA strain contains 20. Since the earliest replicating origin subset across the *S. cerevisiae* genome is only 60 origins in a population of cells (Alvino et al., 2007; Yabuki et al., 2002), 18 additional early-replicating rDNA origins in a single cell could easily generate significant competition for limiting replication factors or nucleotides in early S phase (Mantiero et al., 2011; Shyian et al., 2016) and cause the observed genome replication delays.

### The increased DNA damage sensitivity in strains with reduced rDNA arises from a synthetic interaction with *fob1Δ*

The genome replication delays in the minimal rDNA strain might contribute to the strain’s increased sensitivity to DNA damage; however, a previous study reported that the additional, untranscribed rDNA copies protect cells from DNA damage by facilitating recombinational repair (Ide 2010). We sought to disentangle the effects of early rDNA replication and DNA repair on DNA damage sensitivity in the minimal rDNA strain by considering two other mutants that replicate rDNA early: *rif1Δ* and *sir2Δ* (Figure 5).

**Figure 5:**
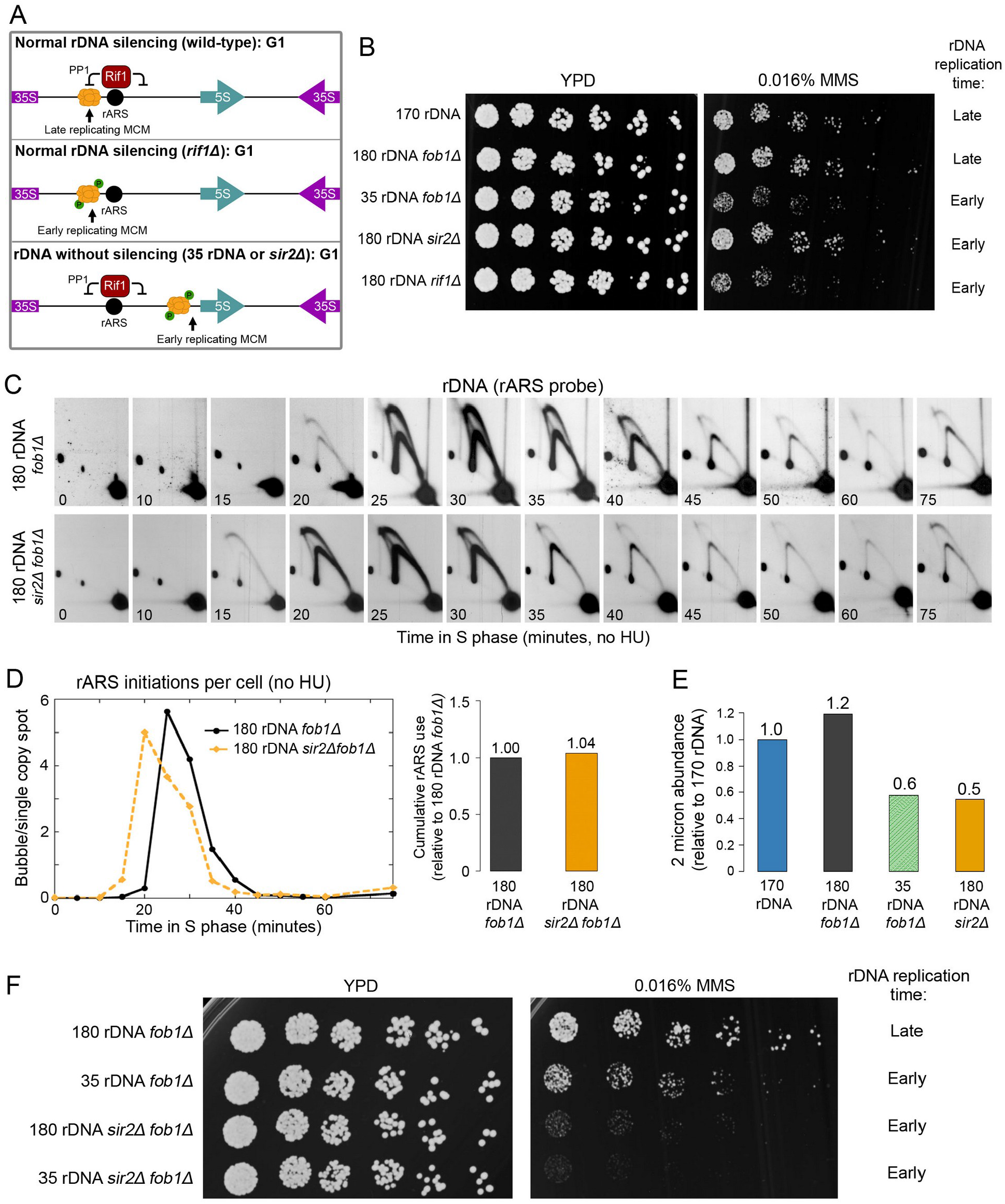
Sensitivity to MMS is due to a synthetic interaction between Fob1p and early rDNA replication. (A) Proposed mechanisms leading to early replicating rDNA. (B) The 35 rDNA *fob1Δ* strain displays increased sensitivity to 0.016% MMS whereas the *sir2Δ* strain and *fob1Δ* strain show wild type sensitivity. (C) Timed 2D gels of rDNA replication initiation in a 170 rDNA *sir2Δ fob1Δ* strain. (D) Quantification of rDNA initiation signal per cell (relative to the 24.4 kb single copy rDNA spot) over the course of S phase for 180 rDNA *fob1Δ*, 35 rDNA *fob1Δ*, and 180 rDNA *sir2Δ fob1Δ* strains. (E) Abundance of the 2 micron plasmid, a parasitic element partially dependent on the *S. cerevisiae* replication machinery, was assessed in logarithmically growing cells using quantitative Southern blotting and normalized to a single copy genomic probe (ChrXV:810). (F) Deletion of *FOB1* increases MMS sensitivity of strains with early replicating rDNA.

Rif1p inhibits local replication initiation by recruiting protein phosphatase 1 (PP1, Glc7p), which prevents the premature activation of the replication helicase subunit Mcm4p (Davé et al., 2014, p. 1; Hiraga et al., 2014; Mattarocci et al., 2014). Because Rif1p binds at the rARS (Hafner et al., 2018; Shyian et al., 2016), wild type rDNA arrays replicate early in Rif1p’s absence. Sir2p is a histone deacetylase responsible for silencing rDNA repeats. Without Sir2p, reduced nucleosome occupancy and increased transcription at active rDNA repeats facilitate the translocation of MCM helicases away from repressive chromatin environments (Foss et al., 2019), likely generated by rARS-bound Rif1p. These unregulated MCM helicases drive early replication initiation at the wild type-length rDNA arrays in *sir2Δ* mutants (Foss et al., 2017; Saka et al., 2013; Yoshida et al., 2014). The minimal rDNA strain shares a critical feature with a *sir2Δ* mutant: its rDNA array is also euchromatic and highly transcribed (French et al., 2003; Ide et al., 2010), likely allowing loaded MCM helicases to translocate away from the rARS-bound Rif1p.

Although the rDNA shifts to early replication in all three backgrounds – minimal rDNA, *sir2Δ*, and *rif1Δ – sir2Δ* mutants did not show increased DNA damage (Figure 6B). This finding seemed to exclude genome replication defects as a sole source of DNA damage sensitivity because these defects were reported to be more severe in *sir2Δ* mutants than in strains with reduced rDNA arrays (Foss et al., 2017; Yoshida et al., 2014). We first confirmed that loss of *SIR2* results in early replicating rDNA (Figure 6C), although we observed no obvious increase in cumulative rDNA origin initiation across S phase (Figure 6D). Second, as a proxy for plasmid maintenance, we examined abundance of the 2 micron plasmid, whose numbers decrease in the presence of replication defects (Maiti and Sinha, 1992; Storici et al., 1995). Both strains with early replicating rDNA (*sir2Δ* and minimal rDNA) showed reduced 2 micron plasmid abundance, to approximately 50% of the levels in wild type control strains (Figure 6E).

**Figure 6:**
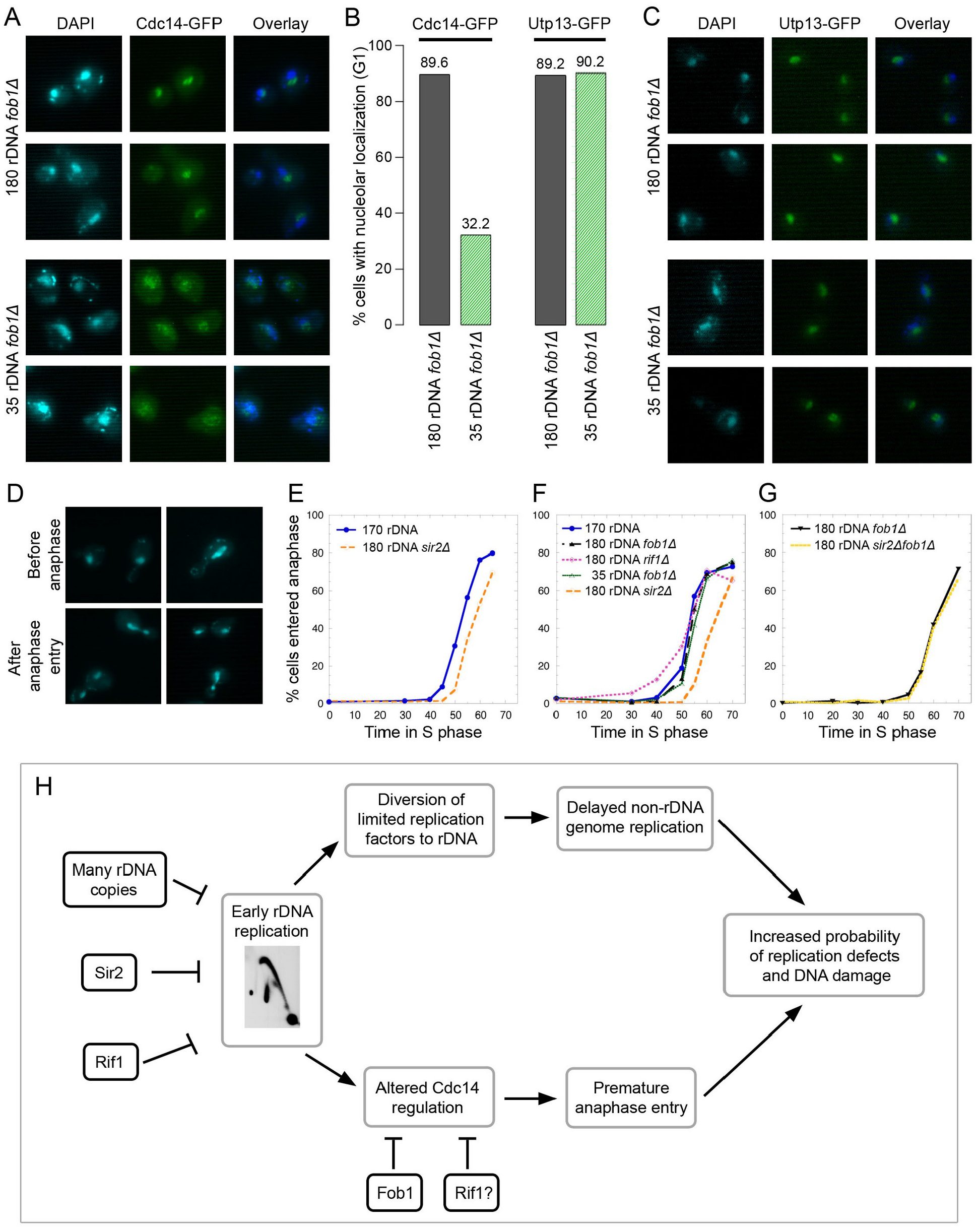
Minimal rDNA strains show poor nucleolar sequestration of Cdc14p and premature anaphase entry. (A) C-terminally tagged Cdc14-GFP was visualized in 180 rDNA *fob1Δ* and 35 rDNA *fob1Δ* cells. (B) Quantification of G1 cells with nucleolar localization of Cdc14-GFP or Utp13-GFP. (C) Utp13-GFP, a nucleolar protein that is involved in rRNA processing, was used to ascertain nucleolar structure (Woolford and Baserga 2013). Both 35 rDNA *fob1Δ* and 180 rDNA *fob1Δ* cells show normal nucleolar structure. (D) Examples of DAPI-stained nuclei representing cells scored as either being “before anaphase” or “after anaphase entry.” (E-G) Quantification of percentage of cells that had entered anaphase for each strain over time. (H) Proposed model regarding consequences of rDNA copy number reduction and early rDNA replication.

Because Sir2p plays a role in restricting genome-wide DNA replication (Hoggard et al., 2020, 2018), we wondered if deletion of *SIR2* could rescue the minimal rDNA strain’s sensitivity to DNA damage and replication stress. We therefore examined MMS sensitivity upon *sir2Δ* deletion in the minimal rDNA strain and the wild type rDNA strain. To our surprise, deletion of *SIR2* in a *fob1Δ* background led to increased MMS sensitivity regardless of rDNA copy number (Figure 6F). This *fob1Δ -*dependent MMS sensitivity was reproducible among multiple strain isolates, each of which was assessed for rDNA copy number (Figure S6A-B), confirming a synthetic defect in DNA damage sensitivity between *fob1Δ* and *sir2Δ*.

Although the rDNA shifts to early replication in all three backgrounds – minimal rDNA, *sir2Δ*, and *rif1Δ – sir2Δ* mutants did not show increased DNA damage (Figure 5B). This finding seemed to exclude genome replication defects as a source of DNA damage sensitivity because these defects were reported to be more severe in *sir2Δ* mutants than in strains with reduced rDNA arrays (Foss et al., 2017; Yoshida et al., 2014). We first confirmed that loss of *SIR2* results in early replicating rDNA (Figure 5C), although we observed no obvious increase in cumulative rDNA origin initiation across S phase (Figure 5D). Second, as a proxy for plasmid maintenance, we examined abundance of the 2 micron plasmid, whose numbers decrease in the presence of replication defects (Maiti and Sinha, 1992; Storici et al., 1995). Both strains with early replicating rDNA (*sir2Δ* and minimal rDNA) showed reduced 2 micron plasmid abundance, to approximately 50% of the levels in wild type control strains (Figure 5E).

Because Sir2p plays a role in restricting genome-wide DNA replication (Hoggard et al., 2020, 2018), we wondered if deletion of *SIR2* could rescue the minimal rDNA strain’s sensitivity to DNA damage and replication stress. We therefore examined MMS sensitivity upon *sir2Δ* deletion in the minimal rDNA strain and the wild type rDNA strain. To our surprise, deletion of *SIR2* in a *fob1Δ* background led to increased MMS sensitivity regardless of rDNA copy number (Figure 5F). This *fob1Δ -*dependent MMS sensitivity was reproducible among multiple strain isolates, each of which was assessed for rDNA copy number (Figure S6A-B), confirming a synthetic defect in DNA damage sensitivity between *fob1Δ* and *sir2Δ*.

We considered that *fob1Δ* might drive sensitivity of strains with early rDNA replication to DNA damage/replication stress. The *fob1Δ* mutation is required to prevent expansion of reduced rDNA arrays (Kobayashi et al., 1998), making it challenging to assess the consequences of the minimal rDNA array in a wild type background. However, we hoped that the rDNA expansion after *FOB1* restoration might be slow enough to capture and assess DNA damage sensitivity of a short rDNA array. To this end, we examined spores from a cross of the minimal rDNA strain (35 rDNA copies; *fob1Δ*) strain with a *MAT*α 150 rDNA *FOB1* strain. Each spore was allowed to form a colony, which was immediately inoculated into culture, and both rDNA copy number and MMS sensitivity were assessed from the same culture. The isolated *FOB1* and *fob1Δ* cells with short rDNA had approximately 55 rDNA copies when plated on MMS (Figure S6C). As suitable controls, we employed the previously engineered *fob1Δ* strains with 55 and 80 rDNA copies, with the former exhibiting greater sensitivity to MMS than the latter. The *FOB1* cells with 55 rDNA copies were more resistant to MMS than *fob1Δ* cells with 55 rDNA copies or minimal rDNA; there was no discernible difference between *FOB1* and *fob1Δ* cells with 150 rDNA copies (Figure S6D). Thus, it is the loss of *FOB1* in combination with rDNA reduction and/or early rDNA replication that causes sensitivity to DNA damage. We therefore propose that the presence of Fob1p itself plays a significant role in the mitigation of early rDNA replication effects, outside of the genome replication delays we observed and the previously reported DNA damage sensitivity (Ide et al., 2010).

### The mitotic exit regulator Cdc14 is mislocalized in the minimal rDNA strain

The replication fork-block protein Fob1p has other roles in addition to its eponymous FOrk-Blocking activity (Krawczyk et al., 2014; Salim et al., 2021; Ward et al., 2000), including sequestration of the mitotic-exit phosphatase Cdc14p (Huang and Moazed, 2003; Stegmeier et al., 2004). We focused on Cdc14p sequestration since delayed genome replication could alter mitotic exit. Cdc14p is recruited to two regions of the rDNA: at the Replication Fork Barrier (RFB) where it is bound to Fob1p and upstream of the 35S transcription start site where it is bound by a yet unidentified factor (Figure S7A; Huang et al., 2006; Huang and Moazed, 2003; Stegmeier et al., 2004). Cdc14p is sequestered in the nucleolus until its release to the nucleus during early anaphase and to the cytoplasm during late anaphase and mitotic exit (Mohl et al., 2009; Shou et al., 1999; Stegmeier et al., 2004; Visintin et al., 1999).

We were curious to determine whether Cdc14p localization would be altered by rDNA copy number reduction in a *fob1Δ* mutant background. We examined localization of Cdc14p-GFP in comparison to DAPI-stained nuclei in both the minimal rDNA and wild type rDNA strains arrested in G1. Almost 90% of wild type rDNA cells showed Cdc14p-GFP normally sequestered to the nucleolus, which excludes DAPI (Figure 6A-B); the remaining cells with Cdc14p-GFP overlapping the DAPI-stained nucleus are likely a consequence of their nucleoli being above or below the nucleus during microscopy. In the minimal rDNA strain, a mere 32.2% of G1 cells showed nucleolar Cdc14p-GFP, with 67.8% of cells showing diffuse nuclear Cdc14p-GFP.

This aberrant Cdc14p localization is not due to a loss of nucleolar integrity. We examined nucleolar structure in our strains using a GFP fusion of Utp13p, a nucleolar protein involved in ribosome biogenesis (Huh et al., 2003; Woolford and Baserga, 2013). Utp13p-GFP localization was identical in both the minimal rDNA and the wild type rDNA strains (Figure 6B-C), indicating that nucleolar structure was not altered, consistent with previous findings (Dauban et al., 2019). The minimal rDNA strain’s aberrant Cdc14p-GFP localization is consistent with a mitotic exit defect that exacerbates effects from delayed genome replication.

### A redundant mitotic exit checkpoint in yeast?

If Cdc14p is involved in coordinating genome replication and mitotic exit, we would expect delayed anaphase entry to accommodate the genome replication defects in strains with early rDNA replication. Given the substantial delays in DNA replication and cyclin signaling, we anticipated late anaphase entry in all strains with early replicating rDNA: the 35 rDNA *fob1Δ* strain, and the *sir2Δ* and *rif1Δ* single mutants. We examined DAPI-stained nuclear morphology across S phase as a proxy for entry into anaphase ((Hartwell et al., 1974; Yellman and Roeder, 2015), Figure 6D). As anticipated, the *sir2Δ* mutant entered anaphase later than the wild type (Figure 6E-F). However, both the minimal rDNA strain and the *rif1Δ* strain entered anaphase at the same time or earlier than their wild type control strains, suggesting that these mutants have lost the mechanism to delay anaphase in response to delayed genome replication (Figure 6F).

The link between genome replication status and anaphase entry appears to be partially dependent on *FOB1*: no difference was observed in anaphase entry between *sir2Δ* and *SIR2* strains in the absence of *FOB1* (Figure 6G). The observed disconnect between S phase completion and anaphase entry mirrors our earlier results with the MMS sensitivity assay: the minimal rDNA, *rif1Δ* and *sir2Δ fob1Δ* strains showed anaphase progression in spite of replication delays as well as increased MMS sensitivity (Figure 5B, F). The *FOB1*-dependent coordination between anaphase entry and genome replication completion suggests that Fob1p acts as a redundant, yeast-specific checkpoint for S phase completion functioning in concert with Cdc14p sequestration (Figure S7).

## DISCUSSION

### Minimal rDNA arrays replicate early and delay genome replication

Here, we characterized the replication consequences of reducing the *S. cerevisiae* rDNA locus from its wild type ∼180 copies to a minimal array of 35 copies. We found altered replication timing not just at the rDNA locus but also across the genome. The minimal rDNA array replicates at the very earliest part of S-phase whereas the full-length rDNA is among the latest replicating regions. This shift in replication timing is explained by the ∼20 additional rDNA initiations that occur in the minimal rDNA array in early S phase. We propose that these additional early-replicating rDNA origins divert the limiting factors required to activate non-rDNA origins, thereby creating delays in replication elsewhere in the genome. Our findings contradict previous interpretations of similar data, whose authors concluded that rDNA copy number reduction leads to replication defects only at the rDNA locus and not in the rest of the genome (Ide et al., 2010). However, replication of other large chromosomes appears visibly impaired in their manuscript (Ide et al. 2010, Figure 2D), consistent with our findings and interpretation.

The dramatic shift of rDNA replication to the early part of S-phase in the minimal rDNA strains comes at a price: impaired plasmid maintenance, heightened sensitivity to HU and MMS, and delayed genome replication. It appears paradoxical that reducing rDNA copy number generates replication stress given that the full-length wild type rDNA array is dramatically shortened in many conditions of imposed replication stress, either in the presence of DNA replication mutants or limiting conditions like HU (Ide et al., 2007; Lynch et al., 2019; Salim et al., 2017; Sanchez et al., 2017). Indeed, rDNA copy number reduction has been proposed to be a compensatory mechanism for replication stress (Ide et al., 2007; Kwan et al., 2013; Salim et al., 2017). We argue that rDNA copy number reduction will affect genome replication only if the rDNA replication timing is affected. This argument is supported by earlier studies describing a mutant that induces replication stress and reduces rDNA copy number (Lynch et al., 2019). The reduced rDNA array in this mutant remained late-replicating, possibly because its copy number reduction was not severe enough to alter replication timing. Further work is necessary to determine the copy number reduction leading to early rDNA replication and whether this shift in replication time is gradual or precipitous.

### A synthetic interaction between early rDNA replication and *FOB1* explains sensitivity to DNA damage in strains with reduced rDNA copy number

Early replicating rDNA causes genome replication defects (this study, (Foss et al., 2017; Ide et al., 2010; Shyian et al., 2016; Yoshida et al., 2014)), yet the increased DNA-damage sensitivity in strains with early-replicating rDNA is not solely a response to delayed genome-wide replication. As we show, *sir2Δ* single mutants have early replicating rDNA with wild type copy number and suffer genome-wide replication defects (Foss et al., 2017) but this background is not sensitive to DNA damage. Comparing the *sir2Δ fob1Δ* and *sir2Δ* strains, we identified a synthetic interaction between *fob1Δ* and early rDNA replication, which results in increased sensitivity to the DNA-alkylating agent MMS.

Previous studies proposed that reduced rDNA copy number sensitizes strains to DNA damage because DNA repair is impaired in short arrays in which all copies are transcribed (Ide et al., 2010). This interpretation was based on the observation that a deletion of the PolI-subunit *RPA135*, and consequently cessation of endogenous rDNA transcription, erased the difference in MMS sensitivity between a 20-copy rDNA strain and a strain purportedly carrying 110 copies. However, deletions of rDNA transcription machinery, including *RPA135*, result in an 80% reduction in rDNA copy number (Brewer et al., 1992; Kobayashi et al., 1998). Thus, the similarity in MMS sensitivity between the two strains could simply stem from similarly low rDNA copy number, which was not verified in the study (Ide et al., 2010).

Moreover, the MMS-sensitive phenotype of the *sir2Δ fob1Δ* strain, which contains over 150 rDNA copies, is inconsistent with the model proposed by Ide *et al*. This strain should have sufficient DNA repair capacity because loss of *SIR2* only moderately increases the number of accessible, presumably actively transcribed rDNA copies (*i.e*. 40% accessible vs 60% non-accessible in wild type yeast; 50% accessible vs 50% non-accessible in *sir2Δ*, (Smith and Boeke, 1997)). However, in our hands, the *sir2Δ fob1Δ* strain shows possibly even greater MMS sensitivity than the minimal rDNA *fob1Δ* strain (Figure 6E). Taken together, our results demonstrate that the capacity for DNA damage repair provided by additional, silenced rDNA copies does not explain the sensitivity of reduced rDNA strains to DNA damage.

### The synthetic interaction between early rDNA replication and *FOB1* uncovers a putative cell cycle checkpoint in yeast

Considering prior evidence and our findings, we propose a model explaining 1) how rDNA replication might coordinate whole genome replication and anaphase entry and 2) how Fob1p might function as a redundant checkpoint in Cdc14 sequestration and mitotic exit.

In a wild type yeast cell (Figure S7A), ∼2000 Cdc14p molecules (Cherry et al., 2012; Ho et al., 2018; Kulak et al., 2014) are sequestered at two sites within an rDNA repeat: at the transcription start site (TSS) and the replication fork barrier (RFB) bound by Fob1p. Cdc14p remains bound at all 180 rDNA repeats during G1 and early S phase. In late S phase, 36 rDNA origins fire and replication forks begin to dislodge Cdc14p from the nearby TSS, but Cdc14p remains bound to Fob1p at the 180 RFB sites. Fob1p, which enforces unidirectional replication (Brewer and Fangman, 1988; Linskens and Huberman, 1988), acts as second, more stable tether for Cdc14p because the RFB can only be replicated by an oncoming fork from an adjacent active origin. Given that a 1 in 5 rDNA origins initiate in a wild type cell (Brewer and Fangman, 1988; Linskens and Huberman, 1988), the nearest active origin is ∼45 kb (5 rDNA arrays) away on average, and it will take the replication machinery ∼30 minutes to reach the stalled fork. Hence, rDNA will complete replication 30 minutes after rDNA origins fire in S phase, consistent with the rDNA locus as the very last region to complete replication. Thus, Cdc14p remains sequestered in the very last region of the genome to replicate, to be fully dislodged by replication at the very end of S phase. We posit that full replication of the rDNA signals the genome-wide replication completion and enables subsequent anaphase entry through Cdc14p release.

In a *fob1Δ* mutant with wild type rDNA copy number (Figure S7B), Cdc14p is not bound to the RFB, but remains associated with the rDNA TSS until rDNA replication completion in late S/early anaphase (Stegmeier et al., 2004). The wild type-length rDNA is still replicated in late S phase and the genome finishes replication by the time of complete Cdc14p release, maintaining coordination of both replication processes. The *fob1Δ* strain retains wild type sensitivity to DNA damage and replication stress agents.

In a *fob1Δ* mutant with minimal rDNA copy number (Figure S7C), Cdc14p nucleolar localization is severely disrupted even prior to S phase. The 35 rDNA copies do not provide enough binding sites for Cdc14p, in particular because Fob1p is missing at the RFB. The minimal rDNA array replicates early, which dislodges Cdc14p fully in early to mid S phase.

Free, unsequestered Cdc14p l will induce anaphase entry while the genome has not yet completed replication. This disconnect between completion of genome replication and anaphase entry may lead to premature cell cycle progression without allowing sufficient time for DNA repair, resulting in increased sensitivity to DNA damage and replication stress agents (Figure 2B, 6A).

In a *sir2Δ* mutant with wild type rDNA copy number (Figure S7D), the rDNA replicates early while replication delays occur elsewhere. This strain shows wild type sensitivity to DNA damage (MMS, Figure 6A). The presence of Fob1p promotes the retention of Cdc14p at the RFB and also enforces slower, unidirectional rDNA replication. Although the rDNA array replicates early, it would still take ∼30 minutes for rDNA replication to be complete. During this time, Cdc14p will remain bound to the RFB via Fob1p while the genome will complete replication, apparently even with rDNA-induced delays. In this way, anaphase entry remains coordinated with genome replication completion. Consistent with this interpretation, we observed delayed anaphase in *sir2Δ* single mutant strains, which suggest anaphase entry remains coordinated with genome replication.

In a *sir2Δ fob1Δ* mutant with wild type rDNA copy number (Figure S7E), Cdc14p is sufficiently bound to the rDNA array in G1, but is missing at RFBs because of the absence of Fob1p. Fob1p absence also leads to more rapid, bidirectional replication of the rDNA array, which is completed in 15 minutes in this strain. This early and rapid replication of the rDNA array will dislodge Cdc14p from the nucleolus while the rest of the genome is in the midst of replicating. Therefore, the *sir2Δ fob1Δ* strain loses coordination between completion of genome replication and anaphase entry, resulting in increased sensitivity to DNA damage (Figure 6E, 7G).

Taken together, the evolutionarily conserved excess of rDNA copies in concert with their late replication act as checkpoint for whole genome replication via Cdc14p sequestration in the nucleolus. Cdc14p sequestration redundantly requires the presence of the yeast-specific Fob1p at RFBs.

### rDNA copy number and genome replication: implications for disease

rDNA copy number can affect the essential processes of whole genome replication and cell cycle progression, extending the phenotypic impact of this genomic element far beyond ribosome biogenesis (Figure 7H). Although *S. cerevisiae* populations typically maintain strain-specific rDNA copy number (Kwan et al., 2016), the repetitive nature of rDNA arrays can allow for rare array contraction below the range of natural variation. Thus, rDNA copy number should be taken into consideration as a background variable when interpreting the consequences of other genetic variants. In fact, rDNA copy number changes are frequently observed after standard *S. cerevisiae* genetic manipulation practices (Kwan et al., 2016).

For metazoans, rDNA copy number reduction may have implications for health outcomes. A recent study reports that rDNA copy number reduction precedes pathogenesis in an mTOR-activated cancer mouse model, suggesting that rDNA reductions may act as driver mutations in certain cancers (Wang and Lemos, 2017; Xu et al., 2017). Hutchinson-Gilford progeroid cell lines show bloated nucleoli (Buchwalter and Hetzer, 2017) and increased DNA damage that appears late in S phase (Chojnowski et al., 2020), echoing phenotypes observed in yeast strains with early replicating rDNA. Although the replication fork barrier protein Fob1p itself is not conserved in metazoans, some Cdc14p homologs are known to localize to the nucleolus (Berdougo et al., 2008; Kaiser et al., 2004; Manzano-López and Monje-Casas, 2020; Saito et al., 2004; Wu et al., 2008). Cdc14p regulation in metazoans may be more sensitive to early rDNA replication without the redundant Fob1p tether.

### How plausible is DNA replication as a gauge of whole genome replication status?

While the nucleolus was originally deemed an oddly mundane location for such an exciting molecule, Cdc14p sequestration in the nucleolus is now recognized as an important hallmark of cell cycle regulation (Amon, 2008). We argue that this mundane organelle, and Cdc14p’s localization within in it, are ideal for cell cycle control. Every cell contains nucleoli, membrane-free organelles that form around the rDNA arrays. Nucleoli and rDNA transcription are highly responsive to cell and organismal physiology, including nutritional status, stress, and aging. The rDNA is late-replicating and associated with Cdc14p whose release into the nucleus coincides with anaphase entry (Figure 7H). The late replication of the rDNA locus may be conserved across a wide variety of species in order to both mitigate replication competition and coordinate replication status with cell cycle progression.

## Acknowledgements

This work was supported by the following funding sources: University of Washington Genome Training Grant T32HG000035 to KLL, NIGMS R35 GM122497 to BJB and MKR, NIGMS R01 GM122088 to C.Q., and NIH grant RM1 HG010461 to C.Q. We are grateful to the Kaeberlein Lab for generously providing us with the Cdc14-GFP and Utp13-GFP strains used to generate strains for this paper. We would like to thank to Dr. Elizabeth Morton, Ashley Hall, Dr. Matthew Crane, Mitsuhiro Tsuchiya, and Benjamin Blue for lively rDNA club discussions, critiques, and moral support. We greatly appreciate Dr. Kerry Bubb for her statistics expertise and help with linear regression. We would also like to extend thanks for the western blotting advice given by Kate Sitko, Barbara Taskinen, and Jason Stephany from the Fowler Lab.

## Author contributions

Conceptualization, E.X.K., G.M.A., B.J.B., C.Q., and M.K.R; Methodology, E.X.K., G.M.A., B.J.B., and M.K.R.; Investigation, E.X.K., G.M.A., K.L.L., P.F.L., H.M.A., X.S.W., S.A.J., J.C.S., M.A.M., M.C., S.B.L., and M.N.; Resources, E.X.K., G.M.A., B.J.B., and M.K.R.; Writing - Original Draft, E.X.K, G.M.A., B.J.B., C.Q., and M.K.R.; Writing - Review & Editing, E.X.K., G.M.A., K.L.L., B.J.B., C.Q., and M.K.R.; Supervision: B.J.B., J.T.C., C.Q., and M.K.R.; Funding Acquisition, B.J.B., M.K.R., and C.Q.

## Declaration of interests

The authors declare no competing interests.

## MATERIALS AND METHODS

### Yeast strains, probe fragments, plasmids, and media

Yeast strains used are listed in Supplementary Table 1. Yeast strains were grown, unless noted otherwise, in synthetic complete media buffered with 1% succinic acid (per liter: 1.45g yeast nitrogen base, 20 g glucose, 10 g succinic acid, 6 g NaOH, 5 g (NH_4_)_2_SO_4_, 2.8 g amino acid powder mix with pH adjusted to 5.8). In cases when YPD medium is used, per liter: 20 g bacto peptone, 10 g yeast extract, and 20 g glucose.

Since we were concerned about possible *de novo* rDNA copy number changes over the course of this study, we froze multiple samples from log-phase cultures that had been CHEF gel verified (Figure 1B), and used these frozen stocks as inoculants for each experiment.

### rDNA reduction

S288c *fob1Δ* strains transformed with the pRDN1-Hyg plasmid, were first isolated by selection for uracil prototrophy, then plated onto medium containing hygromycin B to select for rDNA copy number reduction (Chernoff et al., 1994; Kobayashi et al., 2001; Kwan et al., 2013). Individual colonies were picked for screening by CHEF gel electrophoresis to measure rDNA copy number (Figure S1). We identified and isolated strains with 35, 45, and 55 copies of rDNA and decided to focus on strains with 35 rDNA copies (“35 rDNA *fob1Δ*” and “35 rDNA^RM^ *fob1Δ”*), restoring endogenous *URA3* to facilitate downstream replication assays. Genetic crosses were used generate prototrophic strains and GFP-tagged strains.

During the isolation of strains with reduced rDNA copy number by this pRDN1-HYG plasmid method, we noticed that 20-25% of the isolates with rDNA reductions had either diploidized or tetraploidized (Figure S1), something we had not previously observed when constructing strains by transformation. Subsequently, we verified ploidy of each strain for each experiment by flow cytometry and used only confirmed haploid strains for each experiment. This frequent increase in ploidy may be related to rDNA reduction by this pRDN1-Hyg method. While interesting, we have not identified the biological mechanism involved and strongly suggest verifying ploidy when this rDNA reduction method is employed in the future.

### Preparation of DNA in agarose plugs

DNA was isolated in agarose plugs according to previously published protocols (Tsuchiyama et al., 2013). Each 90 μL plug contained either ∼10^8^ stationary phase cells for CHEF gels, ∼5 x 10^7^ log phase cells for rDNA 2D gels, or ∼10^8^ log phase cells for single-copy origin 2D gels. Collected cells were washed with 50 mM EDTA, resuspended in 90 μL 0.5% SeaPlaque GTG agarose in 50 mM EDTA, and transferred into plug molds. Once solidified, plugs were incubated in 1 mL spheroplasting solution (1 M sorbitol, 20 mM EDTA, 10 mM Tris-HCl pH7.5, 14 mM β-mercaptoethanol, 0.5 mg/mL Zymolyase-20T (Amsbio)) for 2-5 hours at 37°C. Plugs were washed once with LDS (1 % lithium dodecyl sulfate, 100 mM EDTA, 10 mM Tris–HCl pH 8.0) and incubated overnight at 37°C in LDS overnight with gentle shaking. Plugs were then washed 3 x 30 minutes in 0.2X NDS (1X NDS pH 9.5: 0.5 M EDTA, 10 mM Tris base, 1% Sarkosyl) and 5 x 30 minutes in TE pH 8.0. Processed plugs were stored at 4°C in TE pH 8.0 until use.

### CHEF gel analysis

We used contour-clamped homogeneous electric field (CHEF) gel electrophoresis to resolve intact *S. cerevisiae* chromosomes. A slice of each genomic DNA agarose plug was embedded in a 0.8% agarose gel (0.5X TBE) and each gel contained one wildtype sample as reference. For most CHEF gels, we ran the samples in 2.3L of 0.5X TBE using a Bio-Rad CHEF-DRII electrophoresis cell at 100V for 66 hours (switch time = 300 to 900 seconds). The gels were then stained with ethidium bromide to visualize all chromosomes, including the rDNA-containing chromosome XII. To examine the size of the excised rDNA array, genomic DNA samples in plugs were digested with BamHI or FspI and then run on a 0.8% CHEF gel at 165 V for 64 hours (switch time = 47 to 170 seconds). Chromosome XII size and rDNA copy number were further examined via Southern blotting. For size comparison, known standards (*H. wingei* and/or Yeast Ladder from New England BioLabs) were included in each CHEF gel run.

### Sample collection for chromosome replication completion assay

Cells were grown to mid-logarithmic phase (2.5 x 10^6^ cell/mL), arrested in G1 with 3 μM α-factor, and released into S phase (by the addition of 0.15 mg/mL Pronase (EMD Millipore)) in the presence of 0.008% MMS. Samples were collected every 20 minutes for CHEF gel electrophoresis and prepared as described above in agarose plugs for CHEF gel electrophoresis. The same Southern blot membrane was probed for all measured chromosomes except for the FspI-excised rDNA.

### Southern blotting

Each gel was transferred to a GeneScreen Hybridization membrane using standard Southern blotting protocols (Tsuchiyama et al., 2013) We then hybridized each sequence of interest using a ^32^P-labeled probe. The blots were exposed to X-ray film and to Bio-Rad Molecular Imaging FX phosphor screens for visualization and quantification of signal intensity. Phosphor screens were scanned using a Bio-Rad Personal Molecular Imaging scanner and analyzed using Bio-Rad’s Quantity One software. Southern blots were often stripped and re-probed with a different sequence of interest (CHEF gel blots, 2D gel blots, density transfer blots). To strip a Southern blot, blots were subjected to two washes of 20 minutes each in 500 mL stripping buffer (0.1% SSC; 1% SDS) that had been heated to 100°C. Blot stripping efficacy was gauged by exposure and quantification of phosphor screens before the next probe hybridization.

### rRNA quantification

rRNA quantification was performed as described (Sanchez et al., 2017). Asynchronous logarithmic phase cells were collected and nucleic acids (RNA and DNA) were isolated using a “Smash & Grab” phenol:chloroform extraction protocol (Radford, 1991). The RNA northern blot was hybridized to a ^32^P-labeled probe for the 25S rRNA sequence. To assess loading normalization, the DNA Southern blot portion was hybridized to a ^32^P-labeled probe for *ACT1*, a single copy gene. The rRNA and *ACT1* blots were separately exposed to S Bio-Rad phosphor screens and *25S* rRNA and *ACT1* DNA intensity was quantified using a Bio-Rad Personal Molecular Imager and Bio-Rad Quantity One software.

### Density transfer

The density transfer protocol was adapted from (Alvino et al., 2007) Dense medium composition was 0.5% ^13^C-labeled glucose, 0.5% ^15^(NH_4_)_2_SO_4_, 0.00145% yeast nitrogen base (YNB), and 1% succinic acid (isotopically-light medium was the same composition with normal glucose and (NH_4_)_2_SO_4_). Cells were cultured in logarithmic phase for at least 10 generations in dense medium with the growth rate assessed for abnormalities. To collect synchronous S-phase cell samples, cultures of ∼2.5 x 10^8^ cell/mL were arrested with 3 µM α-factor for 1.25 population doublings (approximately 2 hours). Once the cell culture achieved >95% G1 arrest, cells were collected and washed 3 times with isotopically light medium containing α-factor. Cells were resuspended in the original volume of isotopically light medium containing 3 µM α-factor and a 100 mL G1 sample was taken for flow cytometry and DNA analysis. Cells were released from G1 into S phase by the addition of 0.15 mg/mL Pronase (EMD Millipore). 100 mL samples were collected and immediately transferred into vessels containing frozen pellets of 40 mL of 0.1% sodium azide in 0.2 M EDTA. The entire set of timed samples was collected before pelleting cells, taking a small aliquot for flow cytometry, and transferring the rest of the dry pellet to -20°C for storage until DNA isolation. DNA was extracted using a phenol:chloroform “Smash & Grab” protocol (see above) with an additional chloroform cleanup. Isolated DNA was digested overnight with EcoRI and then centrifuged in CsCl to separate replicated from unreplicated DNA. Cesium chloride gradients were drip-fractionated and the collected samples were analyzed using slot blots and hybridization to microarrays.

### Flow cytometry

Cells for flow cytometry were fixed in 70% ethanol before processing for flow cytometry. Fixed cells were washed with 50 mM sodium citrate, sonicated, and resuspended in 500 μL 50 mM sodium citrate. RNase A was added to a concentration of 2.5 mg/mL and the samples were incubated for 1 hour at 50°C. Proteinase K (50 μL of 20 mg/mL) was then added and cells were incubated another hour at 50°C before staining with 1X Sytox Green. Cells were analyzed on a BD Canto II flow cytometer and flow cytometry data was analyzed using FlowJo software.

### 2D gel electrophoresis

Cells from the 180 rDNA *fob1Δ* strain, the 35 rDNA *fob1Δ* strain, and the 180 rDNA *sir2Δ fob1Δ* strain were grown in logarithmic phase to a culture density of ∼2.5 x 10^6^ cells/mL. Cultures were then arrested in α-factor for 1.25 doublings before being released into S phase by addition of Pronase (0.15 mg/mL). Samples were collected every 5 minutes: 100 mL for analysis of single-copy genomic origins or 30 mL for analysis of rDNA origins. Collection vessels contained frozen pellets of 0.1% sodium azide in 0.2 M EDTA to halt growth. Cells were washed once with 50 mM EDTA, a small sample taken for flow cytometry, and the remaining dry cell pellets were stored at -20°C until preparation for 2D gel electrophoresis. To extract DNA for 2D gels, cells were embedded in three 90 μL 0.5% SeaPlaque agarose plugs and prepared as CHEF gel plugs. For each 2D gel, each plug was washed 3 x 20 minutes in the appropriate restriction buffer with 1X BSA (100 μg/mL). The solution was then removed and the DNA was digested for 5 hours by addition of 3 μL restriction enzyme directly onto each plug, and then subjected to standard 2D gel electrophoresis methods (Brewer and Fangman, 1987), Southern blotted and hybridized for the sequence of interest. Cumulative origin initiation was estimated by integrating the area under the curve generated from plotting “rDNA initiations per cell” across time.

For 2D gels of cells in hydroxyurea (HU), 20 mL of the culture was transferred to another flask for the “no HU” control to check that cells would have had normal release into S phase. Hydroxyurea was added to the remaining culture to a final concentration of 200 mM and 10 minutes later, 0.15 mg/mL Pronase was added to both cultures to release the cells into S phase. Samples were collected every 30 minutes and prepared as above.

### Plasmid maintenance assay

Cells that contained *ARS1* plasmids (Kwan *et al*. 2013) were grown to logarithmic phase in selective medium (YPD + 200 μg/mL G418) and then released into non-selective medium (YPD) for the plasmid maintenance assay. Cells were kept in logarithmic phase growth and samples were collected approximately every 4 hours over the course of 48 hours. The growth rate was monitored to ascertain the number of generations/divisions between samples. DNA was extracted from cells using the “Smash & grab” protocol, digested with XmnI, and run on an agarose gel to resolve the 5.4 kb plasmid *ARS1* fragment from the 3.4 kb genomic *ARS1* fragment (used as a “per cell” loading control). The gel was then Southern blotted and the membrane hybridized to a ^32^P-labeled *ARS1* fragment. *ARS1* plasmid abundance was assessed through Southern blotting and normalized to the endogenous *ARS1* locus. We quantified the amount of signal from plasmid *ARS1* and genomic *ARS1* for each sample and generated a plasmid maintenance curve for each strain, from which we were able to calculate the rate of plasmid loss per generation and estimate significance using linear regression.

### Spot assays

Cells were grown to log-phase, diluted in sterile water in 3-fold dilutions, and 2.5 μL was spotted onto YPD plates containing either no drug (control), 0.016% methyl methanesulfonate (MMS), or 200 mM hydroxyurea (HU). Plates were scanned after 40-48 hours of growth at 30°C.

### Cycloheximide sensitivity assay

Cells were grown to log phase, upon which 3 x 10^4^ log-phase cells were transferred to each well in a 96-well plate containing 150 μL medium per well and the appropriate concentration of cycloheximide (0-200 ng/mL). Each condition was performed in triplicate and optical densities were measured at 30°C for 48 hours using a Bio-Tek reader. The maximum log-phase growth rate was manually calculated for each well.

### Western blotting

Cells were grown in log phase to ∼2.5 x 10^8^ cell/mL before α-factor arrest and release. For each strain, 1.5 mL was collected for protein extraction and 1 mL was collected for flow cytometry. Collected cell pellets were resuspended in 200 μL SUMEB buffer (1% SDS, 8 M urea, 10 mM MOPS pH 6.8, 10 mM EDTA, 0.01% bromophenol blue) supplemented with protease inhibitors and 5% β-mercaptoethanol. Glass beads (∼100 μL 0.5 mm acid-washed) were added and cells were vortexed for 3 minutes. Lysates were incubated at 65°C for 10 minutes with intermittent shaking and then centrifuged for 5 minutes at 4°C at 20,000 x g. The clarified supernatant was transferred to a new tube and protein concentration was assessed using a Qubit (Thermo Fisher). For each sample, 15 μg of protein was run on a Novex Tris-acetate SDS-PAGE gel and transferred to a nitrocellulose membrane for immunoblotting. HRP-conjugated antibodies against HA (Sigma Aldrich #12013819001) and Pgk1 (Abcam #ab197960) were used in this work.

### Microscopy

Cells were fixed according the protocol described on the Koshland lab web site (http://mcb.berkeley.edu/labs/koshland/Protocols/MICROSCOPY/gfpfix.html): collected cell pellets were resuspended in paraformaldehyde solution (4% paraformaldehyde, 3.4% sucrose) and incubated at room temperature for 15 minutes. Cells were then washed once with KPO_4_/sorbitol solution and resuspended in 50 μL KPO4/sorbitol solution (0.1 M KPO_4_ pH 7.5, 1.2 M sorbitol) and stored at 4°C until visualization. Before fluorescence microscopy, cells were sonicated and incubated with 0.5 μg/ml DAPI for at least an hour.

## Notes

### Competing Interest Statement

The authors have declared no competing interest.

### Summary of Updates

General figure and text reorganization, title change, significant text edits.

