## Supplemental data for "Coordination of genome replication and anaphase entry by rDNA copy number in *S. cerevisiae*"

#### Kwan Supplementary Table 1: Strain list

| Name | Genotype |
|---|---|
| B150 | S288c MATa ; BY rDNA (170 copies) |
| B30 | S288c MATa *his3Δ0 leu2Δ0 met15Δ0 fob1::cloNAT*; BY rDNA (35 copies) |
| EK342 | S288c MATa *fob1::cloNAT*; BY rDNA (35 copies) |
| EK68 | S288c MATa *fob1::cloNAT*; BY rDNA (180 copies) |
| R90 | S288c MATa; RM rDNA (100 copies) |
| R30 | S288c MATa *his3Δ0 leu2Δ0 lys2Δ0 fob1::cloNAT;* RM rDNA (30 copies) |
| EK040 | S288c MATa *his3Δ0 leu2Δ0 met15Δ0 ura3Δ0 fob1::cloNAT*; BY rDNA (55 copies) |
| EK051 | S288c MATa *his3Δ0 leu2Δ0 met15Δ0 fob1::cloNAT*; BY rDNA (45 copies) |
| EK057 | S288c MATa *his3Δ0 leu2Δ0 lys2Δ0 fob1::cloNAT*; RM rDNA (45 copies) |
| EK100 | S288c MATa *his3Δ0 leu2Δ0 met15Δ0 ura3Δ0 fob1::cloNAT*; BY rDNA (80 copies) |
| EK105 | S288c MATa *ura3::KanMX RAD53-2HA-URA3*; BY rDNA (180 copies) |
| EK109 | S288c MATa *his3Δ0 leu2Δ0 met15Δ0 ura3Δ0 fob1::NAT RAD53-2HA-URA3*; BY rDNA (35 copies) |
| EK125 | S288c MATa *fob1::cloNAT CLB5-3HA-KanMX6*; BY rDNA (180 copies) |
| EK130 | S288c MATa *his3Δ0 leu2Δ0 met15Δ0 CLB5-3HA-KanMX6*; BY rDNA (35 copies) |
| EK139 | S288c MATa *fob1::cloNAT SIC1-3HA-KanMX6*; BY rDNA (35 copies) |
| EK142 | S288c MATa *fob1::cloNAT SIC1-3HA-KanMX6*; BY rDNA (180 copies) |
| EK150 | S288c MATa *sir2::cloNAT HML::KanMX*; BY rDNA (150 copies) |
| EK151 | S288c MATa *sir2::cloNAT HML::KanMX*; BY rDNA (100 copies) |
| EK152 | S288c MATa *his3Δ0 leu2Δ0 met15Δ0 fob1::cloNAT*; BY rDNA (35 copies) + pUC19-KanMX-KwCEN7-ARS1 |
| EK154 | S288c MATa *his3Δ0 leu2Δ0 lys2Δ0 fob1::cloNAT*; RM rDNA (35 copies) + pUC19-KanMX-KwCEN7-ARS1 |
| EK156 | S288c MATa *fob1::cloNAT*; BY rDNA (180 copies) + pUC19-KanMX-KwCEN7-ARS1 |
| EK177 | S288c MATa *his3Δ0 leu2Δ0 met15Δ0 fob1::cloNAT sml1::HYG*; BY rDNA (35 copies) |
| EK182 | S288c MATa *fob1::cloNAT sml1::HYG*; BY rDNA (180 copies) |
| EK184 | S288c MATa; BY rDNA (170 copies) + pUC19-KanMX-KwCEN7-ARS1 |
| EK185 | S288c MATa; RM rDNA (100 copies) + pUC19-KanMX-KwCEN7-ARS1 |
| EK194 | S288c MATa *his3Δ0 leu2Δ0 met15Δ0 ura3Δ0 fob1::cloNAT CDC14-GFP:HIS3*; BY rDNA (35 copies) |
| EK202 | S288c MATa *his3Δ0 leu2Δ0 met15Δ0 ura3Δ0 fob1::cloNAT CDC14-GFP:HIS3*; BY rDNA (170 copies) |
| EK339 | S288c MATα *fob1::NAT*; BY rDNA (35 copies) |
| EK360 | S288c MATa *sir2::cloNAT fob1::cloNAT HML::KanMX*; BY rDNA (180 copies) |
| EK361 | S288c MATa *sir2::cloNAT fob1::cloNAT HML::KanMX*; BY rDNA (170 copies) |
| EK375 | S288c MATa *fob1::cloNAT rif1::KanMX*; BY rDNA (35 copies) |
| EK376 | S288c MATa *fob1::cloNAT sir2::cloNAT HML::KanMX*; BY rDNA (35 copies) |
| EK379 | S288c MATa *fob1::cloNAT rif1::KanMX*; BY rDNA (180 copies) |
| EK396 | S288c MATa *his3Δ0 leu2Δ0 met15Δ0 ura3Δ0 UTP13-GFP:HIS3 fob1::cloNAT*; BY rDNA (35 copies) |
| EK425 | S288c MATa *his3Δ0? leu2Δ0? met15Δ0? ura3Δ0 fob1::NAT UTP13-GFP:HIS3*; BY rDNA (~150 copies) |
| EK465 | S288c MATa *fob1::NAT SLD2-HA:KanMX*; BY rDNA (35 copies) |
| EK466 | S288c MATa *fob1::NAT SLD2-HA:KanMX*; BY rDNA (180 copies) |
| EK468 | S288c MATa *his3Δ0? leu2Δ0? met15Δ0? URA3 rif1::KanMX*; BY rDNA (180 copies) |

**Kwan Table S2: Comparison of $T_{rep}$ (time at half maximal replication, minutes)**

| Genomic region $T_{rep}$ | 180 rDNA *fob1Δ* | 35 rDNA *fob1Δ* |
|---|---|---|
| rDNA | 40.0 | 28.9 |
| *ARS305* | 30.2 | 29.7 |
| *ARS607* | 30.0 | 29.7 |
| *ARS501* | 33.6 | 37.2 |
| *ARS735.5* | 37.5 | 40.0 |
| ChrV:53400 | 36.0 | 37.5 |

### Kwan Figure S1: Characterization of strains generated with rDNA reductions.

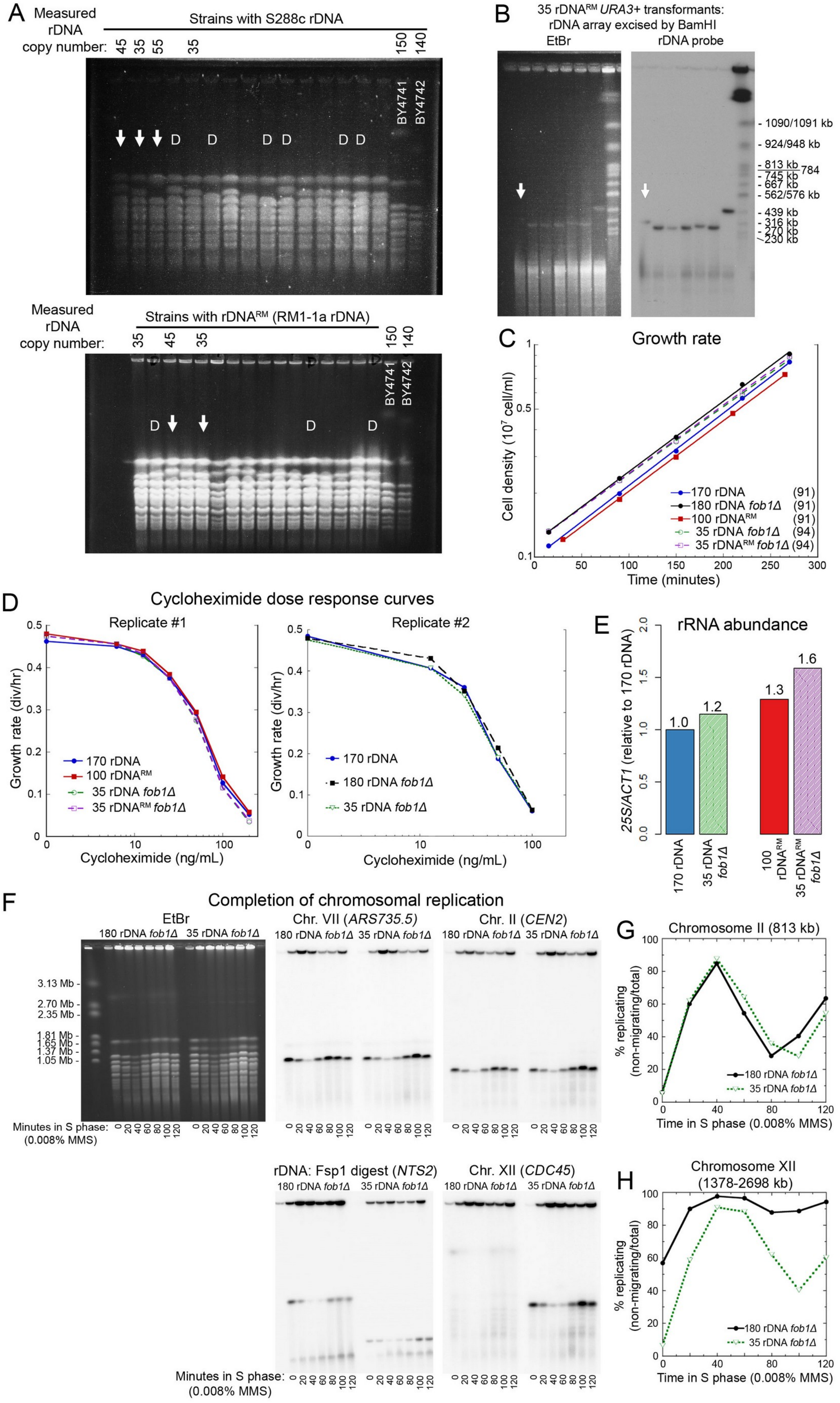

### Kwan Figure S2: Flow cytometry analyses of cell cycle progression.

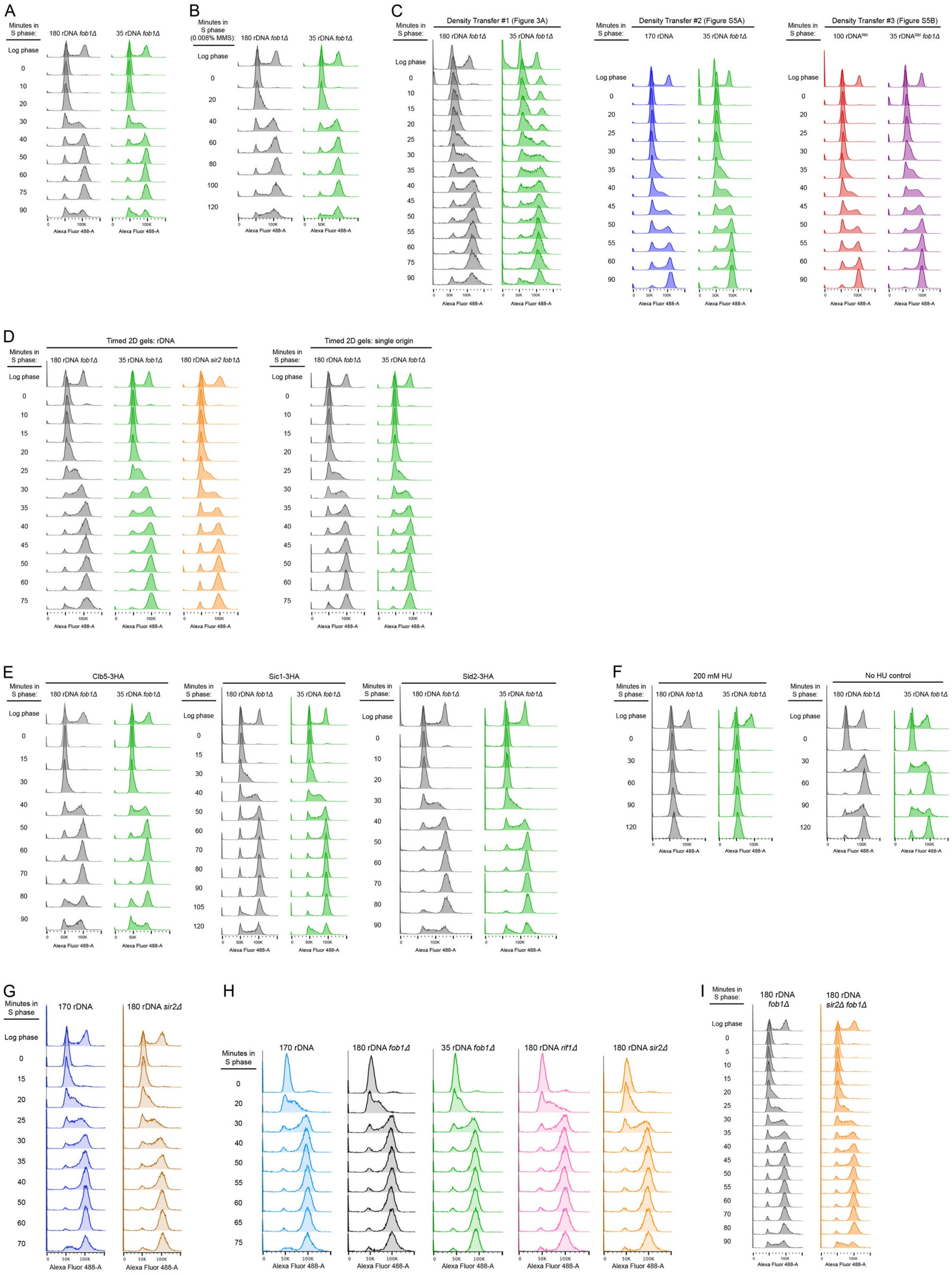

**Kwan Figure S3: Density transfer replication kinetics and timed 2D gels.**

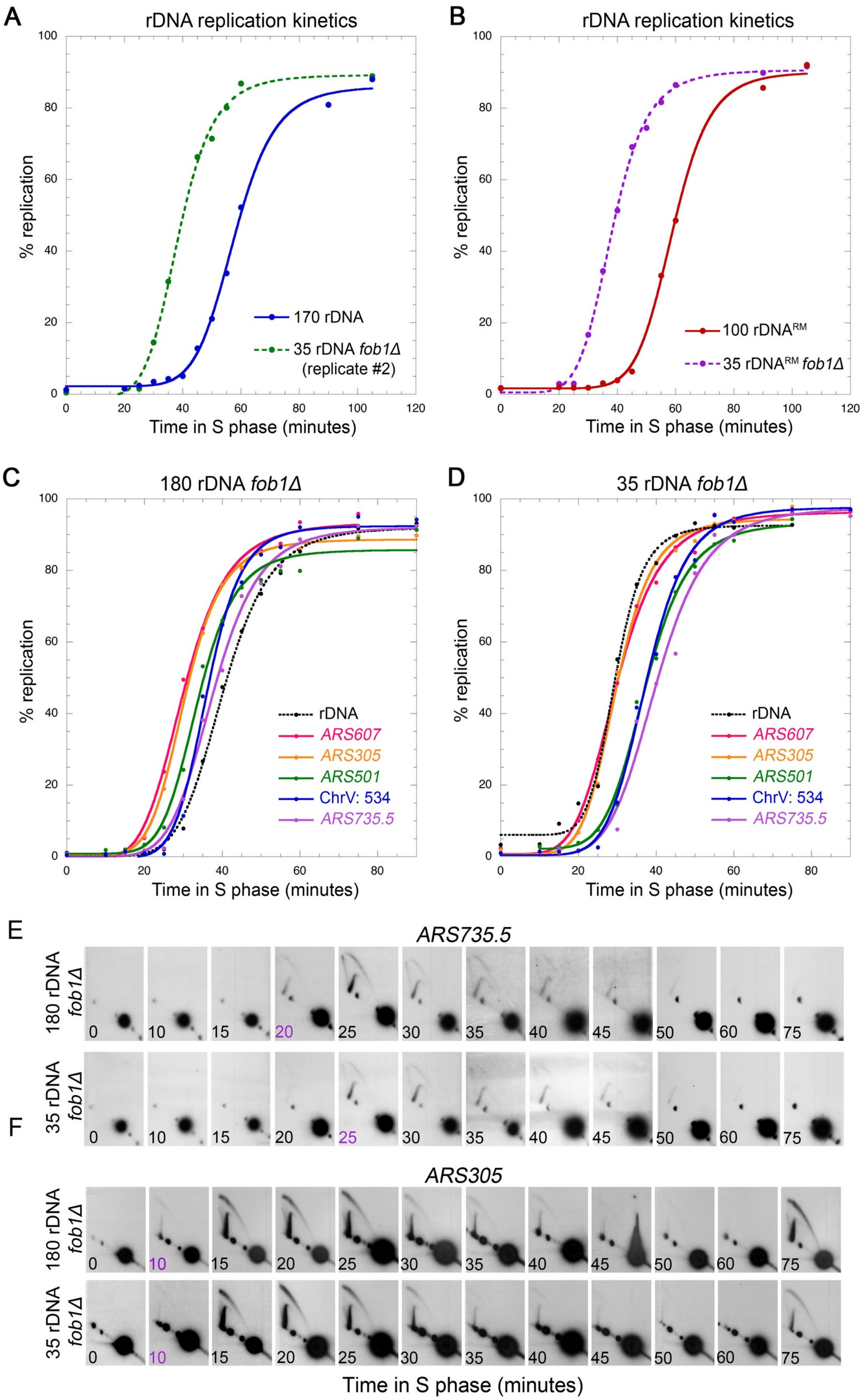

**Kwan Figure S4: Microarray analysis of density transfer samples for all chromosomes.**

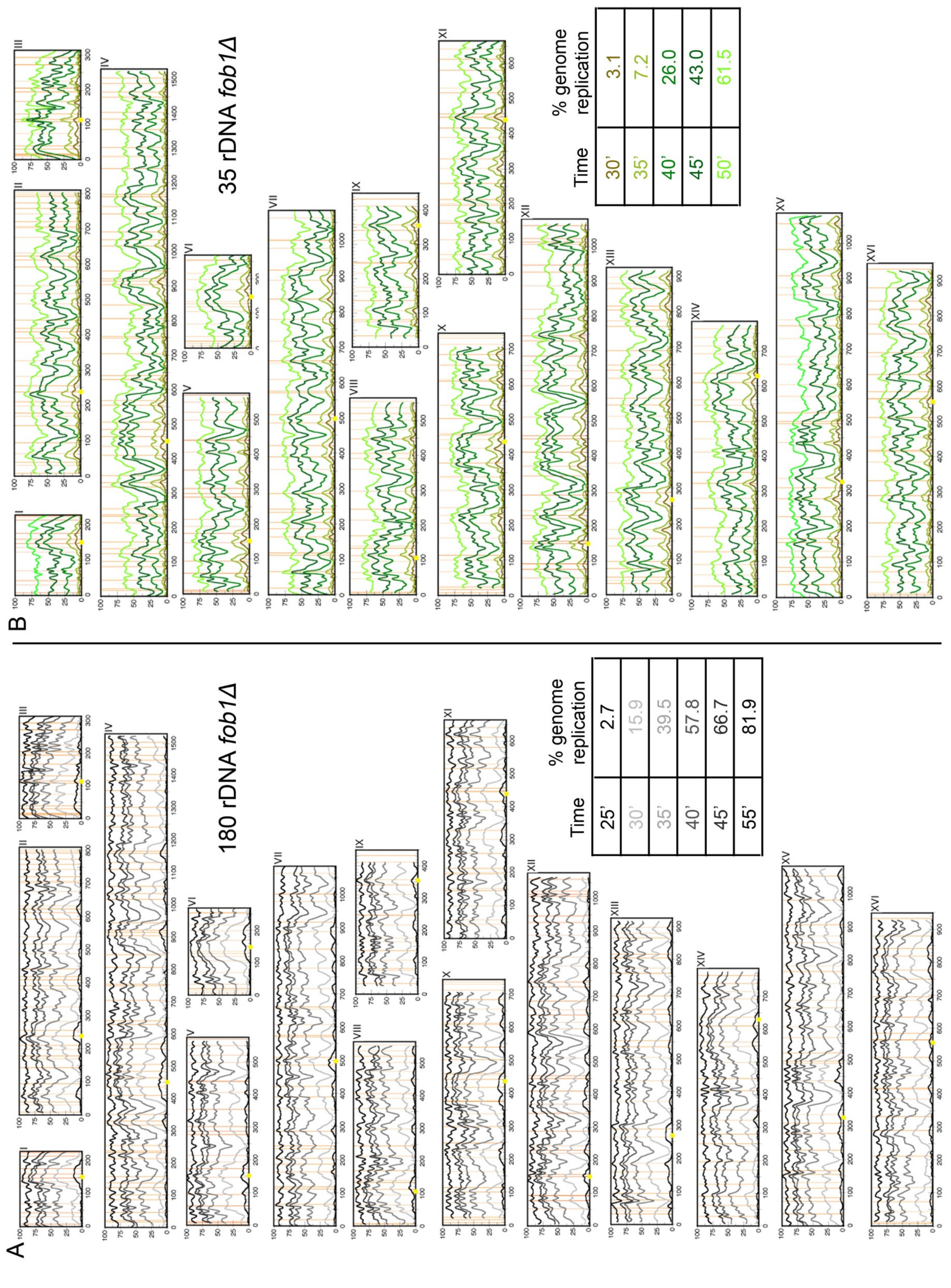

**Kwan Figure S5: Comparison of chromosomal replication profiles at matched replication time or matched % genomic replication. 2D gels used for rDNA quantification during early S phase.**

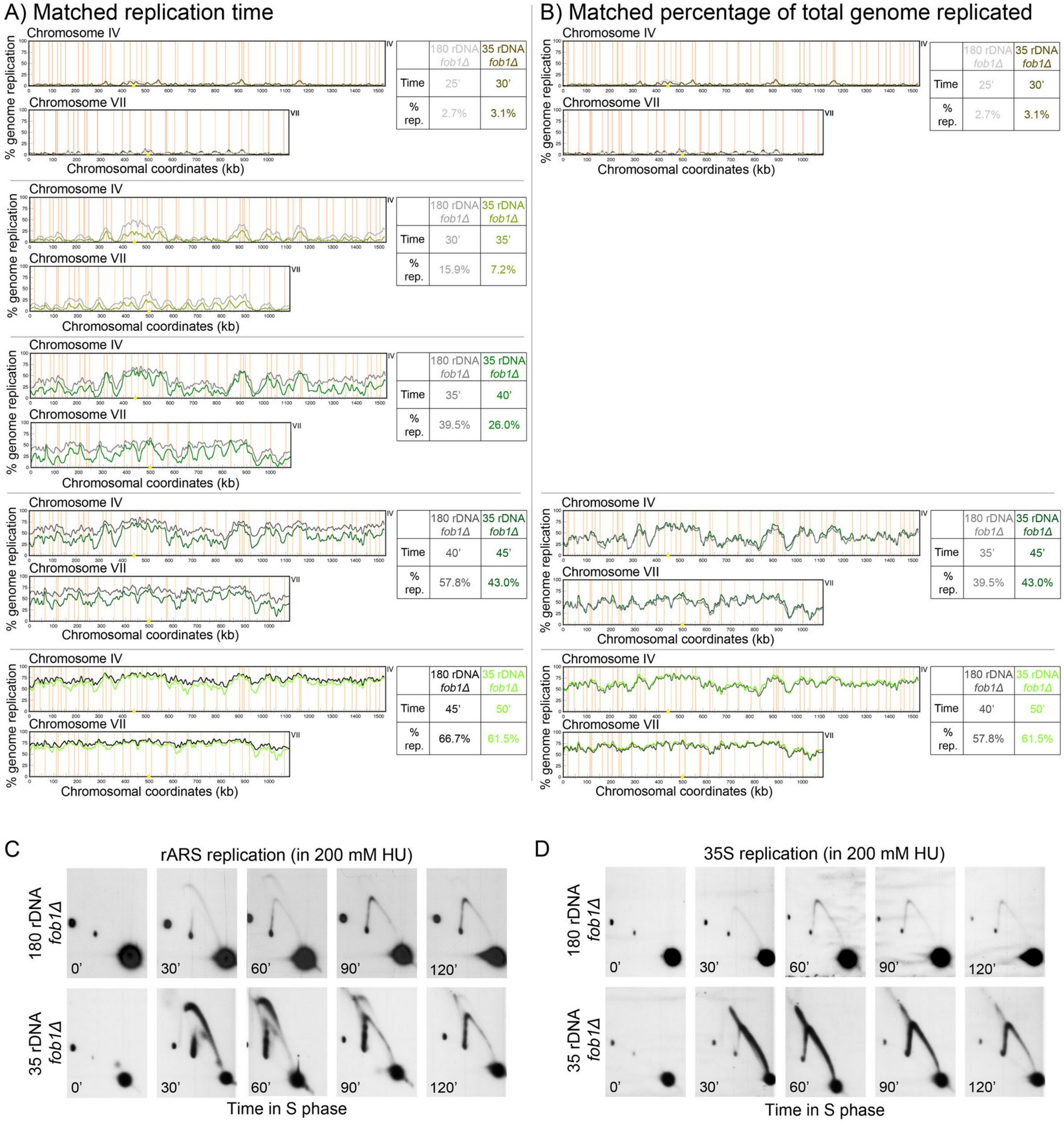

### Kwan Figure S6: CHEF gel of biological replicates examined for MMS sensitivity.

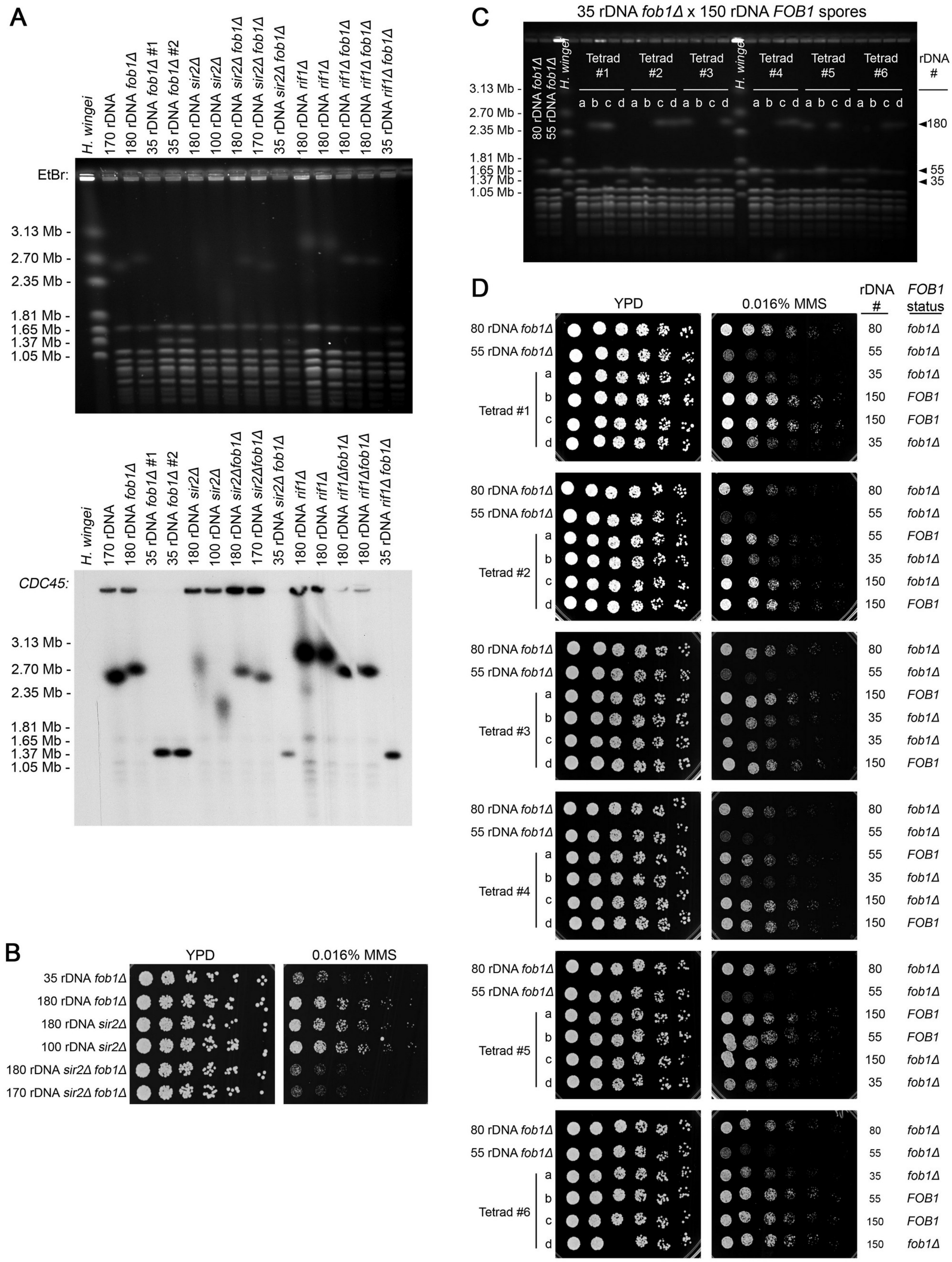

**Kwan Figure S7, model: rDNA replication and Cdc14p sequestration coordinate proper anaphase entry upon completion of whole genome replication**

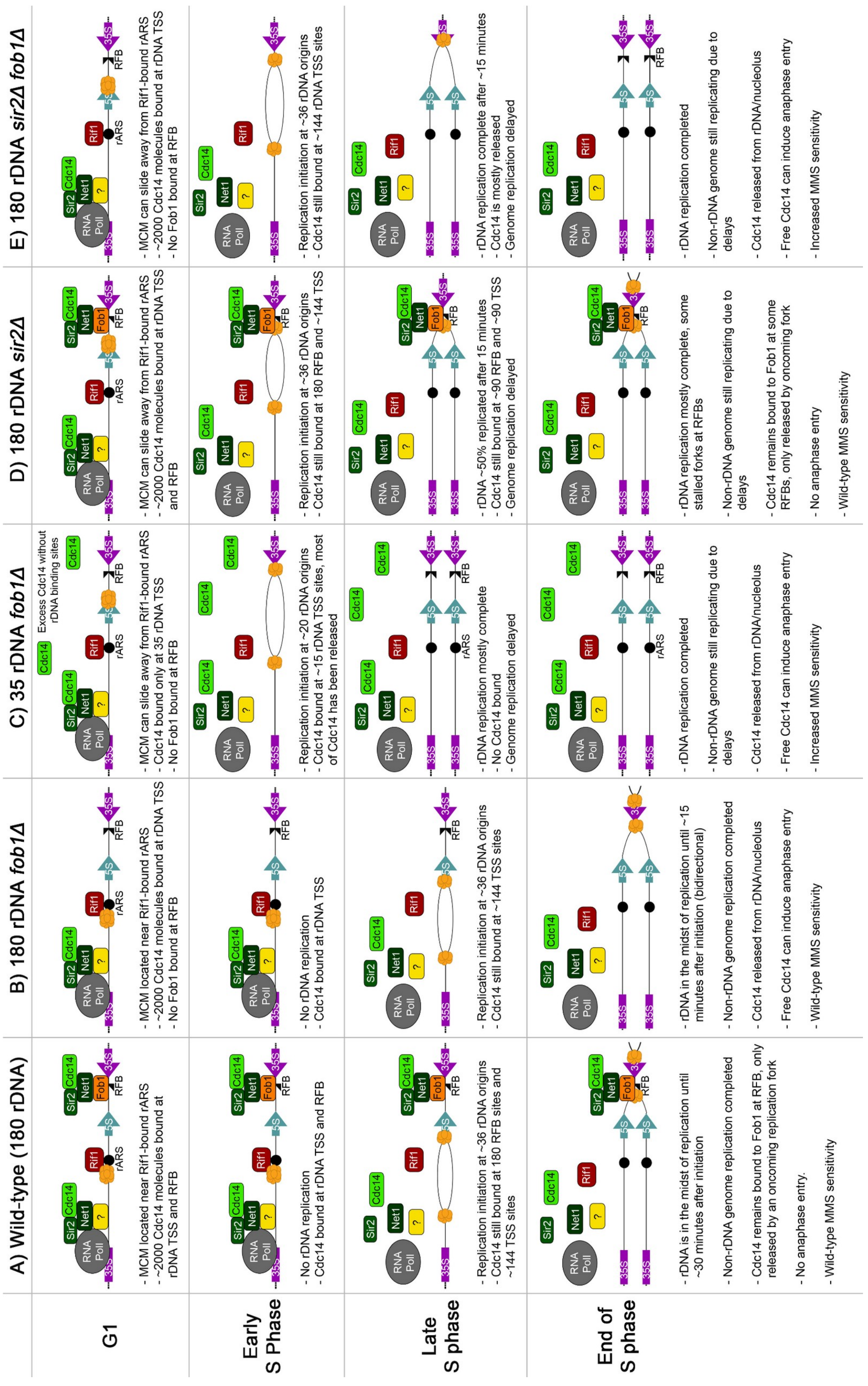

**KWAN SUPPLEMENTAL FIGURE LEGENDS:**

**Figure S1: Characterization of strains generated with rDNA reductions.** We assessed at least 40 isolates after rDNA reduction in *ura3Δ* strains with either S288c rDNA (wild-type) or rDNA with a weakened rARS (RM11-1a rDNA or "$rDNA^{RM}$"). (A) Isolates were initially assessed using chromosome XII size as a proxy for rDNA copy number. Isolates marked with a "D" indicate strains that had diploidized/tetraploidized. Arrows denote strains selected for this study. (B) Example of CHEF gel used to estimate size of a BamHI-excised rDNA array for *URA3+* transformants. We re-introduced *URA3* into strains via a lithium acetate transformation based method. This transformation method can alter rDNA copy number, as seen between the different transformants. Full length *S. cerevisiae* chromosomes from strain BY4741 were used as a ladder. Arrow denotes the 35 $rDNA^{RM}$ *fob1Δ* transformant selected for this study. (C) Growth rate measurements of the five strains. Doubling times (min) indicated in parentheses. (D) Strain sensitivity to the ribosomal translation inhibitor cycloheximide was assessed using a dose response curve in medium containing 0-200 ng/mL cycloheximide. (E) Quantification of rRNA abundance relative to 170 rDNA strain (wild type). Nucleic acid (genomic DNA and total RNA) from each strain was run as a single sample on a gel, which was then split for both northern blotting for 25S rRNA and Southern blotting for a single copy gene (*ACT1*). 25S rRNA abundance was quantified and normalized to the *ACT1* Southern blot loading control. (F) Chromosome replication completion assay: Ethidium bromide stained CHEF gel was blotted and hybridized to radioactive probes for the sequences in Figure 2 and ARS735.5 (Chr. VII), CEN2 (Chr. II), and CDC45 (Chr. XII). The samples from the same CHEF gel plug were digested with FspI, run, and hybridized for NTS2 (rDNA specific probe). (G,H) Quantification of replication completion of Chromosome II and Chromosome XII in this assay.

**Figure S2: Flow cytometry analyses.** Flow cytometry analysis of cell cycle progression from G1 release for (A) S phase, (B) S phase in the presence of 0.008% MMS (Figure 2C-G, S1D), (C) density transfer experiments (Figure 3, S3-5), (D) timed 2D gel analysis (Figure 3C, S3), (E) Western blots for Clb5-3HA/Sic1-3HA/Sld2-3HA (Figure S5), (F) timed 2D gel analysis in 200 mM HU (Figure 4), and (G-I) anaphase entry analysis (Figure 7).

**Figure S3: Density transfer replication kinetics and timed 2D gels.** Comparison of rDNA replication kinetics shown for (A) 170 rDNA and 35 rDNA *fob1Δ* strains and (B) 100 $rDNA^{RM}$ and 35 rDNARM *fob1Δ* strains. The 35 rDNA *fob1Δ* data shown in (A) is a biological replicate of Figure 3A data. Replication kinetics of many genomic $T_{rep}$ values in Figure 3B are plotted together for (C) the

180 rDNA *fob1Δ* strain and (D) the 35 rDNA *fob1Δ* strain. Timed 2D gel electrophoresis: (E) *ARS735.5* is a late-replicating origin and shows delayed replication in the 35 rDNA *fob1Δ* strain. (F) *ARS305* is an early, efficient origin and has similar replication initiation time in the 35 rDNA *fob1Δ* and 180 rDNA *fob1Δ* strains.

**Figure S4: Microarray analysis of density transfer samples for all chromosomes.**

**Figure S5: Comparison of chromosomal replication profiles at matched replication time or matched % genomic replication. 2D gels used for rDNA quantification during early S phase.** (A,B) Comparison of chromosomes IV and VII in the 180 rDNA *fob1Δ* strain (data from Figure 2, S4A) and 35 rDNA *fob1Δ* strain (data from figure S3A, 4B) matched for (A) absolute replication time or (B) total genome replication percentage. Time of replication onset was defined as the time of first perceptible replication signals from both kinetic curve and microarray data (first visible peaks). The genome-wide replication delay in the 35 rDNA *fob1Δ* strain can be seen 5 minutes after replication onset, but comparison of matched genome replication level shows that no region appears to be specifically delayed. (C,D) Full time course of 2D gels examining cells released into S phase in the presence of 200 mM HU, quantified for graphs in Figure 4D,F. 2D gels were probed with both (C) the rARS probe fragment and then (D) the 35S probe.

**Figure S6: CHEF gel of biological replicates examined for MMS sensitivity.** (A) CHEF gel examining chromosome XII size in different strain isolates, stained with ethidium bromide (left) or probed with *CDC45* on chromosome XII (right). (B) MMS sensitivity spot assay replication examining multiple isolates of the *sir2Δ* and *sir2Δ fob1Δ* strains. (C,D) The increased DNA damage sensitivity seen in strains with reduced rDNA copy number is dependent on *fob1Δ.* Strains with reduced rDNA copy number and intact *FOB1* were generated by crossing the 35 rDNA *fob1Δ* strain with a MATα *FOB1* strain with 150 rDNA copies. Tetrad colonies were immediately inoculated, grown to stationary phase, and cells from the same culture were used for both CHEF gel plugs and for MMS spot assays. Previously isolated *fob1Δ* strains with 80 and 55 rDNA copies were included as controls. (C) Ethidium bromide-stained CHEF gel assessing rDNA copy number of all strains. (D) MMS sensitivity spot assay of tetrad spores and control strains. *FOB1* status and estimated rDNA copy number are indicated. Restoration of *FOB1* rescues MMS sensitivity in strains with 55 rDNA copies.
